# Targeting methionine synthase in a fungal pathogen causes a metabolic imbalance that impacts cell energetics, growth and virulence

**DOI:** 10.1101/2020.06.04.131862

**Authors:** Jennifer Scott, Monica Sueiro-Olivares, Benjamin P. Thornton, Rebecca A. Owens, Howbeer Muhamadali, Rachael Fortune-Grant, Darren Thomson, Riba Thomas, Katherine Hollywood, Sean Doyle, Royston Goodacre, Lydia Tabernero, Elaine Bignell, Jorge Amich

**Affiliations:** Manchester Fungal Infection Group (MFIG), Division of Infection, Immunity and Respiratory Medicine, School of Biological Sciences, Faculty of Biology, Medicine and Health, University of Manchester, Manchester Academic Health Science Centre, Manchester, UK; School of Biological Sciences, Faculty of Biology, Medicine and Health, University of Manchester, Manchester Academic Health Science Centre, Manchester, UK; Department of Biology, Maynooth University, Co. Kildare, Ireland; Department of Biochemistry, Institute of Integrative Biology, University of Liverpool, Liverpool, UK; Manchester Institute of Biotechnology. University of Manchester, Manchester, UK

## Abstract

There is an urgent need to develop novel antifungals to tackle the threat fungal pathogens pose to human health. In this work, we have performed a comprehensive characterisation and validation of the promising target methionine synthase (MetH). We uncover that in *Aspergillus fumigatus* the absence of this enzymatic activity triggers a metabolic imbalance that causes a reduction in intracellular ATP, which prevents fungal growth even in the presence of methionine. Interestingly, growth can be recovered in the presence of certain metabolites, which evidences that *metH* is a conditionally essential gene. As this implies that for a correct validation MetH should be targeted in established infections, we have validated the use of the tetOFF genetic model for fungal research and optimised its performance to mimic treatment of established infections. We show that repression of *metH* in growing hyphae halts growth *in vitro*, which translates into a beneficial effect when targeting established infections using this model *in vivo*. Finally, a structural-based virtual screening of methionine synthases reveals key differences between the human and fungal structures and unravels features in the fungal enzyme that can guide the design of novel specific inhibitors. Therefore, methionine synthase is a valuable target for the development of new antifungals.

**IMPORTANCE:** Fungal pathogens are responsible for millions of life-threatening infections on an annual basis worldwide. The current repertoire of antifungal drugs is very limited and, worryingly, resistance has emerged and already become a serious threat to our capacity to treat fungal diseases. The first step to develop new drugs often is to identify molecular targets which inhibition during infection can prevent pathogen growth. However, the current models are not suitable to validate targets in established infections. Here we have characterised the promising antifungal target methionine synthase in great detail, using the prominent fungal pathogen *Aspergillus fumigatus* as a model. We have uncovered the underlying reason for its essentiality and confirmed its druggability. Furthermore, we have optimised the use of a genetic system to show a beneficial effect of targeting methionine synthase in established infections. Therefore, we believe that antifungal drugs to target methionine synthase should be pursued and additionally, we propose that antifungal targets should be validated in a model of established infection.

## INTRODUCTION

Fungal pathogens represent an increasing risk to human health (1), with over one billion people worldwide affected by mycoses annually. Many of these mycoses are superficial infections of the skin, nails or mucosal membranes and although troublesome are usually not life-threatening. However, some fungi cause devastating chronic and invasive fungal infections, which result in an estimated 1.6 million deaths per year (2). Incidences of invasive infections caused by *Aspergillus, Candida, Cryptococcus* and *Pneumocystis* species are increasing (3), a cause for serious concern as these genera are responsible for 90% of deaths caused by mycoses (4). Despite the availability of antifungal drugs, mortality rates for invasive aspergillosis, invasive candidiasis, cryptococcal meningitis and *Pneumocystis jirovecii* pneumonia are intolerably high, reaching over 80%, 40%, 50% and 30% respectively (2, 5). There are currently only four classes of antifungals in clinical use to treat invasive infections (azoles, echinocandins, polyenes and flucytosine), all suffering from pharmacological drawbacks including toxicity, drug-drug interactions and poor bioavailability (6, 7). With the sole exemption of flucytosine, which is only used in combinatory therapy with amphotericin B for cryptococcal meningitis and *Candida* endocarditis (7), the current antifungals target critical components of the fungal cell membrane or cell wall (8), which represents a very limited druggable space. The rise of antifungal resistance presents an additional challenge as mortality rates in patients with resistant isolates can reach 100%, making the development of new antifungal drugs increasingly critical for human health (1, 9). Targeting fungal primary metabolism is broadly considered a valid strategy for the development of novel antifungals, as it is crucial for pathogen virulence and survival (10, 11). A primary example of success of this strategy is olorofim (F901318), a novel class of antifungal that targets the pyrimidine biosynthesis pathway (12), which is currently in clinical trials.

Methionine synthases catalyse the transfer of a methyl group from *N*5-methyl-5,6,7,8-tetrahydrofolate (CH_3_-THF) to L-homocysteine (Hcy). Two unrelated protein families catalyse this reaction: cobalamin dependent methionine synthases (EC 2.1.1.13) and cobalamin independent methionine synthases (EC 2.1.1.14). Members of both families must catalyse the transfer of a low active methyl group from the tertiary amine, CH_3_-THF, to a relatively weak nucleophile, Hcy sulfur. Cobalamin dependent enzymes facilitate this transfer by using cobalamin as an intermediate methyl carrier (13). By contrast, cobalamin independent enzymes directly transfer the methyl group from CH_3_-THF to Hcy (14). Logically, proteins of each family differ significantly both at amino acid sequence (15) and 3D structure level (16).

We have previously shown that the methionine synthase-encoding gene is essential for *A. fumigatus* viability and virulence, which led us to propose it as a promising target for antifungal drug development (17). In support of this, a systematic metabolic network analysis by Kaltdorf and colleagues identified methionine synthase as a promising antifungal drug target worthy of investigation (11). Methionine synthase has also been described as essential for *Candida albicans* viability (18, 19) and necessary for *Cryptococcus neoformans* pathogenicity (20), which suggests that a drug developed against this enzyme may have a broad spectrum of action. Moreover, fungal methionine synthases are cobalamin independent, differing significantly from the cobalamin dependent human protein at the amino acid sequence level: only 11.2% identity, 20.4% similarity and 60.2% gaps when aligned the *A. fumigatus* and human proteins using L-Align from EMBL (21, 22). Therefore, it should be possible to develop drugs with low toxicity potential.

Target validation is critical and has been suggested as the most important step in translating a new potential target into a viable drug target because of its role in achieving efficacy in patients (23). Indeed a retrospective analysis from AstraZeneca’s drug pipeline showed that projects that had performed a more thorough target validation were less likely to fail: 73% of the projects were active or successful in Phase II compared with only 43% of projects without such extra target validation (24). Therefore, in this work we aimed to further substantiate methionine synthase’s potential as an antifungal drug target, before advancing the drug discovery process. In particular, we were interested in 1) unravelling the mechanistic basis of methionine synthase essentiality in *A. fumigatus*, which is needed to fully explore the potential of this enzyme as drug target and to be able to anticipate drug resistance mechanisms; and 2) developing *in vivo* models of infection to mimic treatment against the target in an established infection and using them to validate methionine synthase as an antifungal drug target.

## RESULTS AND DISCUSSION

### Methionine synthase enzymatic activity is essential for Aspergillus fumigatus viability

We had previously demonstrated that the methionine synthase encoding gene is essential for *A. fumigatus* viability and virulence (17); however, the underlying reason for this essentiality was still unclear. To address this question, here we have constructed strains that express the *metH* gene under the control of a tetOFF system recently adapted for *Aspergillus* (25) in two different *A. fumigatus* wild-type backgrounds, ATCC46645 and A1160. The advantage of the tetOFF system over other regulatable systems is that doxycycline (Dox) can be added to downregulate gene expression in growing hyphae (Fig. S1A), and thus this system permits investigation of the consequences of the repression of an essential gene in growing mycelia. The constructed *metH_tetOFF* strains (*H_OFF*) grew as the wild type in the absence of Dox, but as little as 0.5 µg/mL was sufficient to completely prevent colony development on an agar plate even in the presence of methionine (Fig. S1B). This corroborates our previous result that methionine synthase is essential for *A. fumigatus* viability and that its absence does not result in a sheer auxotrophy for methionine (17).

Methionine synthase forms an interjection between the trans-sulfuration pathway and the one carbon metabolic route (Fig. 1A), as the enzyme utilizes 5-methyl-tetrahydrofolate as co-substrate. Therefore, the essentiality of *metH* might be due to required integrities of the trans-sulfuration pathway or of the one carbon metabolic route. Alternatively, it could be that the presence of the enzyme itself is essential, either because its enzymatic activity is required or because it is fulfilling an unrelated additional role, as being part of a multiprotein complex. To start discerning among these possibilities, we constructed a double *ΔmetGΔcysD* mutant, blocked in the previous step of the trans-sulfuration pathway, and a *ΔmetF* deletant, which blocks the previous step of the one-carbon metabolic route (Fig. 1A). As we had previously observed (26), to rescue fully *ΔmetF*’s growth the media had to be supplemented with methionine and other amino acids, as the folate cycle is necessary for the interconversion of serine and glycine and plays a role in histidine and aromatic amino acid metabolism (27, 28). Consequently, we added a mix of all amino acids except cysteine and methionine to the S-free medium for this experiment. Phenotypic tests (Fig. 1B) confirmed that the *ΔmetGΔcysD* and *ΔmetF* mutants were viable and could grow in the presence of methionine. In contrast, the *H_OFF* conditional strain could not grow under restrictive conditions even in the presence of the amino acid mix and methionine (Fig. 1B). Therefore, the MetH protein itself, and not the integrity of the trans-sulfuration and one-carbon pathways, is essential for *A. fumigatus* viability. Interestingly, the methionine auxotroph *ΔmetGΔcysD* was avirulent in a leukopenic model of pulmonary aspergillosis (Fig. S1C), suggesting that the amount of readily available methionine in the lung is very limited, not sufficient to rescue its auxotrophy. Indeed, the level of methionine in human serum was calculated to be as low as ∼20 µM (29, 30), which was described as insufficient to support the growth of various auxotrophic bacterial pathogens (31) and we have also observed that is not enough to rescue growth of the *A. fumigatus ΔmetGΔcysD* auxotroph.

**Figure 1.**
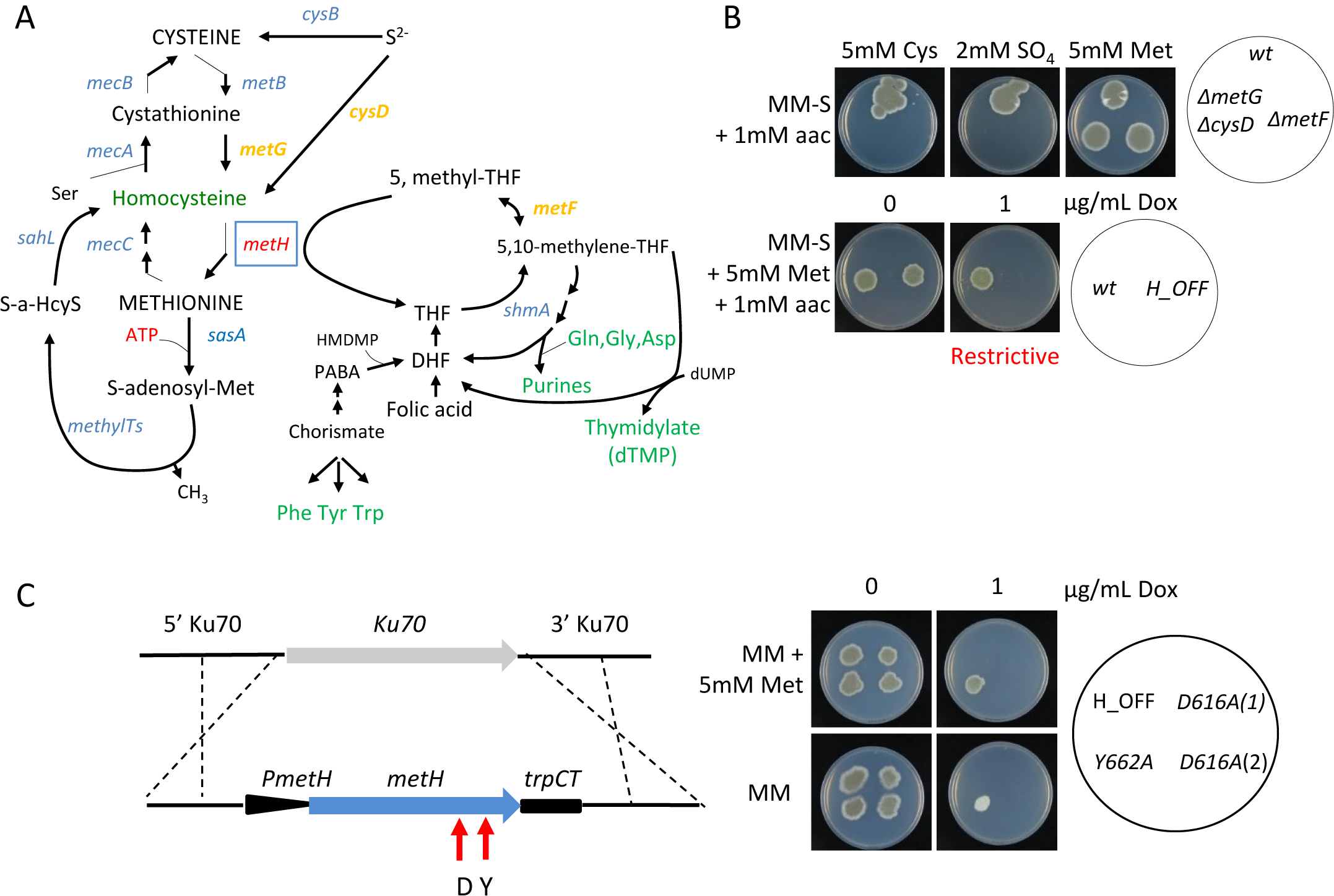
Methionine synthase (MetH) enzymatic activity is essential for *A. fumigatus* viability. **A)** Schematic representation of the trans-sulfuration pathway and its intersection with the one carbon metabolic route. **B)** Both strains, a *ΔmetF* mutant (which blocks the one carbon metabolism route) and a *ΔmetGΔcysD* mutant (which blocks the trans-sulfuration pathway) could grow in the presence of methionine. In contrast, the *metH_tetOFF* strain (*H_OFF*) could not grow in restrictive conditions (+Dox) even if methionine was supplemented. The phenotypic analysis was repeated in three independent experiments. Representative plates are shown. **C)** A second copy of the *metH* gene under the control of its own promoter was introduced in the innocuous *Ku70* locus of the *H_OFF* background strain. Two point-mutated versions of the gene were introduced, one that causes a D→A substitution in amino acid 616 and another one that causes a Y→A substitution in amino acid 662. In non-restrictive conditions all strains were able to grow as the parental *H_OFF* strain. In the presence of Dox and absence of met, the Y662A protein was able to trigger significant growth, suggesting that its enzymatic activity is impaired but not blocked. In the presence of Met the Y662A strain grew as well in restrictive as in non-restrictive conditions, suggesting that partial enzymatic activity is sufficient to cover the essential function of MetH. In restrictive conditions and absence of methionine, the D616A strain was not able to grow, indicating that enzymatic activity is blocked in this mutated protein. The D616A was also not able to grow in the presence of Dox and met, indicating that enzymatic activity is required for the essential function of MetH. The phenotypic analysis was repeated in three independent experiments. Representative plates are shown.

Essentiality of the MetH protein could be directly linked to its enzymatic activity or, alternatively, the protein could be performing an additional independent function. To discern between these two possibilities, we constructed two strains that express single-point mutated versions of MetH from the innocuous Ku70 locus of the *H_OFF* background strain, under the control of its native promoter (Fig. 1C). These point mutations, *metH*^*g2042A>2C*^ (D616A) and *metH*^*g2179TA>9GC*^ (Y662A), were previously described to prevent conformational rearrangements required for activity of the *C. albicans* methionine synthase (32). In the absence of Dox, these strains grew normally, as they expressed both the wild-type MetH, from the tetOFF promoter, and the mutated version of the protein (Fig. 1C). In the presence of Dox, when the wild-type *metH* gene was downregulated, the Y662A strain grew on sulfate worse than in non-restrictive conditions, but still to a significant extent, suggesting that this point mutation did not completely abrogate enzymatic activity (Fig. 1C). Interestingly, Y662 grew normally on methionine, showing that methionine can compensate for a partial reduction of MetH activity (Fig 1C). The D616A mutated protein was confirmed to be stable as a GFP-tagged version of this protein could be visualised in −Dox conditions (Fig. S1D, strain detailed later in the manuscript). Interestingly, the D616A strain (two isolates were tested) was not able to grow on sulfate (Fig. 1C), demonstrating that enzymatic activity was fully blocked. Nor could it grow on methionine (Fig. 1C), indicating that enzymatic activity is required for viability even in the presence of the full protein. All these phenotypes support the conclusion that methionine synthase enzymatic activity is required for viability.

### Absence of methionine synthase enzymatic activity results in a shortage of crucial metabolites, but does not cause toxic accumulation of homocysteine

The absence of methionine synthase enzymatic activity has two direct consequences, which could cause deleterious effects and therefore explain its essentiality (Fig. 1A). It could cause an accumulation of the potentially toxic substrate homocysteine and/or a shortage of the co-product tetrahydrofolate (THF). THF is directly converted to 5,10-methylene-THF, which is required for the synthesis of purines and thymidylate (TMP), and thus for DNA synthesis; additionally, as purine biosynthesis requires Gln, Gly and Asp, and THF *de novo* synthesis requires chorismate (precursor of aromatic amino acids), a shortage of THF might cause a depletion of amino acids (Fig. 1A). To investigate if the depletion of any of these metabolites underlies MetH essentiality, we supplemented the media with a number of precursors and potentially depleted metabolites (Fig. 2A). Added as sole supplement, only adenine was able to trigger growth, but to a minimal degree. Further addition of a mixture of all amino acids noticeably improved growth. Supplementation with adenine and guanine (purine bases) did also reconstitute noticeable growth, which was not enhanced with further addition of amino acids. Folic acid was also capable of reconstituting growth, but only when amino acids were added as the sole N-source (Fig. S3). However, no combination of compounds was able to reconstitute growth to the wild-type level. This suggests that a shortage of relevant metabolites derived from THF, prominently adenine, partially accounts for methionine synthase essentiality, but cannot explain it completely. In other fungi, as *Pichia pastoris* (33) or *Schizosaccharomyces pombe* (34), summplementation with methionine and adenine was found to restore growth of a *metH* mutant to wild-type levels, denoting them as combined auxotrophs. In *A. fumigatus* it seems to be more complex because supplementation with methionine, adenine and other amino acids could still not fully restore growth, suggesting that more factors are implicated.

**Figure 2.**
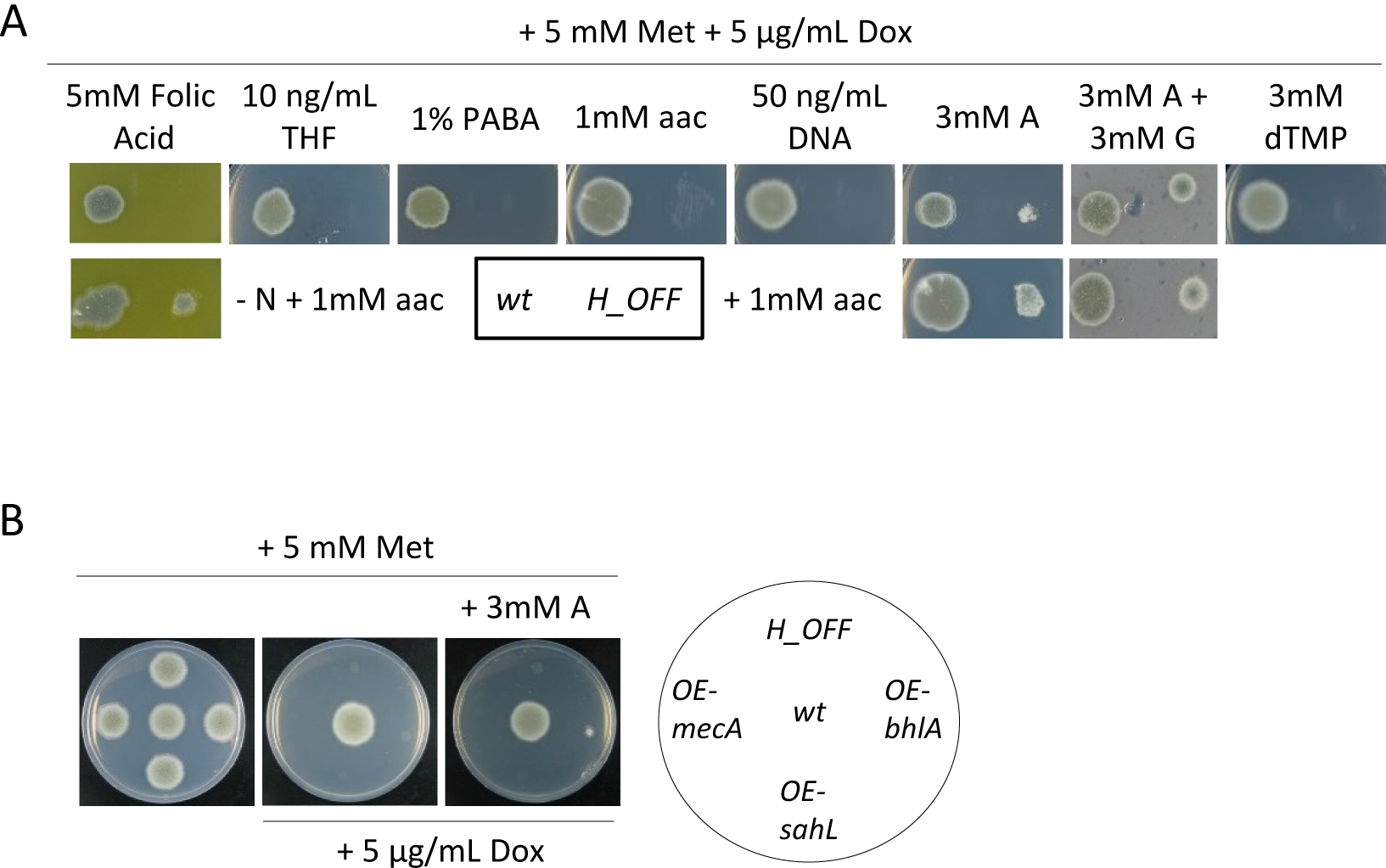
Shortage of important downstream metabolites, but not toxic accumulation of homocysteine, partially accounts for MetH essentiality. **A)** Supplementation of the growth media with a variety of downstream metabolites (Fig. 1A) showed that purines (adenine A and guanine G, 3 mM) could slightly reconstitute growth of the *H_OFF* strain in restrictive conditions. Further supplementation of amino acids improves growth, but not to the wild-type levels. Folic acid (5 mM) could also reconstitute growth when amino acids are the only N-source. **B)** Overexpression of genes that could detoxify a potential accumulation of homocysteine did not reconstitute growth in the absence of MetH activity. Further addition of adenine did not improve growth. The phenotypic analyses were repeated in three independent experiments. Representative plates are shown.

To investigate if homocysteine could be accumulating to toxic levels in the absence of MetH activity, we over-expressed several genes that should alleviate its accumulation. To this aim we designed and constructed the plasmid pJA49, which allows direct integration of any ORF to episomally overexpress genes in *A. fumigatus*. Plasmid pJA49 carries the *A. nidulans* AMA1 autologous replicating sequence (35, 36) and the hygromycin B resistance gene (*hygrB*) as a selection marker. A unique *Stu*I restriction site allows introduction of any PCR amplified ORF in frame under the control of the *A. fumigatus* strong promoter *hspA* (37) and the *A. nidulans trpC* terminator (Fig S2A). Using this plasmid, we produced a strain in the *H_OFF* background that episomally overexpresses *mecA*, encoding cystathionine-β-synthase, which converts homocysteine to cystathionine (Fig. 1A). Homocysteine exerts toxic effects through its conversion to *S*-adenosylhomocysteine, which causes DNA hypomethylation (38, 39), or to homocysteine thiolactone, which causes *N*-homocysteinylation at the ε-amino group of protein lysine residues (40, 41). Consequently, we also constructed strains that episomally over-express genes that could detoxify those products: the *S*-adenosyl-homocysteinase lyase encoding gene *sahL* (AFUA_1G10130) or the *A. nidulans* homocysteine thiolactone hydrolase encoding gene *blhA* (AN6399) (*A. fumigatus* genome does not encode any orthologue) (Fig. S2B). However, despite a strong over-expression of the genes (Fig. S2C&D), none of them could rescue growth of the *H_OFF* strain in restrictive conditions (Fig. 2B). Therefore, our over-expression experiments suggest that homocysteine accumulation is not responsible for *metH* essentiality, but additional experiments such as quantification homocysteine levels, which are currently challenging, would be required to further support this hypothesis. Addition of adenine to the medium did not improve growth of the overexpression strains further than that of the *H_OFF* background (Fig. 2B), indicating that methionine synthase essentiality seemingly is not a combined effect of homocysteine accumulation and depletion of THF-derived metabolites. Toxic accumulation of homocysteine was speculated to be the underlying reason of methionine synthase essentiality in both *Candida albicans* and *Cryptococcus neoformans* (19, 20) but our results suggest that this is not the case in *A. fumigatus*. Therefore, we propose that the previous assumption should be revisited in other fungal pathogens.

### Methionine synthase repression triggers a metabolic imbalance that causes a decrease in cell energetics

Aiming to identify any adverse metabolic shift in the absence of MetH and/or accumulation of toxic compounds that could explain its necessity for proper growth, we performed a metabolomics analysis, via gas chromatography-mass spectrometry (GC-MS), comparing the metabolites present in wild-type and *H_OFF* strains before and 6 h after Dox addition. Before Dox addition both strains clustered closely together in a Principal Component Analysis (PCA) scores plot (Fig. S4A), showing that their metabolic profiles are highly similar. However, 6 h after Dox addition the strains clusters became clearly separated, denoting differential metabolite content. Analysis of the differentially accumulated metabolites (full list can be consulted in Table S1) using the online platforms MBRole (42) and Metaboanalyst (43, 44) did not reveal any obvious metabolic switch, probably due to the rather small number of metabolites that could be identified by cross-referencing with the Golm library (http://gmd.mpimp-golm.mpg.de/). Manual inspection of the metabolites pointed out interesting aspects. Firstly, the methionine levels were not significantly different, which demonstrates that methionine supplementation in the growth medium triggers correct intracellular levels in the *H_OFF* strain; this undoubtedly rules out that a shortage of methionine could be the cause of the essentiality of methionine synthase. Secondly, we detected a significantly lower amount of adenosine in the *H_OFF* strain compared with the wild-type after Dox addition (Fig. 3A), which is in agreement with our previous result that supplementation of adenine can partially reconstitute growth in the absence of MetH. We did not find accumulation of compounds with a clear toxic potential upon *metH* repression. Nevertheless, we detected a lower amount of several amino acids (Phe, Ser, Glu, Pro, Ile, Thr, Ala and Asp, Fig. S4B), which suggests that the cells may enter into growth arrest upon *metH* repression. Interestingly, we noticed a significantly lower accumulation of some metabolites of the glycolysis pathway and TCA cycle (Fig. 3A) and some other mono and poly-saccharides (Fig. S4B). These variations could reflect a low energetic status of the cells upon *metH* repression. Indeed, we found that the level of ATP significantly decreased in the *H_OFF* strain, but not in the wild-type, upon Dox addition (Fig. 3B). Therefore, we evaluated if supplementation of the medium with substrates that have the potential to increase cell energetics can rescue *H_OFF* growth in restrictive conditions. We found that when pyruvate, which can directly be converted to acetyl-CoA to enter the TCA cycle, was added as the sole carbon source *H_OFF* growth was reconstituted in restrictive conditions to the same level as the wild-type (Fig. 3C). Growth was limited for both strains, as pyruvate does not appear to be a good carbon source (45). However, the presence of glucose in the medium precluded the reconstitution of growth of *H_OFF* (Fig. S3), as it has been described to prevent pyruvate uptake in *S. cerevisiae* (46). We next tested the capacity of ATP to be used an alternative energy source and to reconstitute growth. To diversify the presence of permeases in the cell membrane, and thus maximise the chance of ATP uptake, we assayed two different N-sources: ammonium (NH_4_^+^, preferred source) and amino acids (Fig. S3). Indeed, when amino acids were the only N-source, supplementation of the medium with ATP reconstituted *H_OFF* growth in restrictive conditions to wild type levels (Fig. 3C). This agrees with the recent observation that eukaryotic cells can uptake ATP and exploit it as an energy source (47). In conclusion, a decrease in cell energetics developed in the absence of methionine synthase seems to explain MetH essentiality for growth.

**Figure 3.**
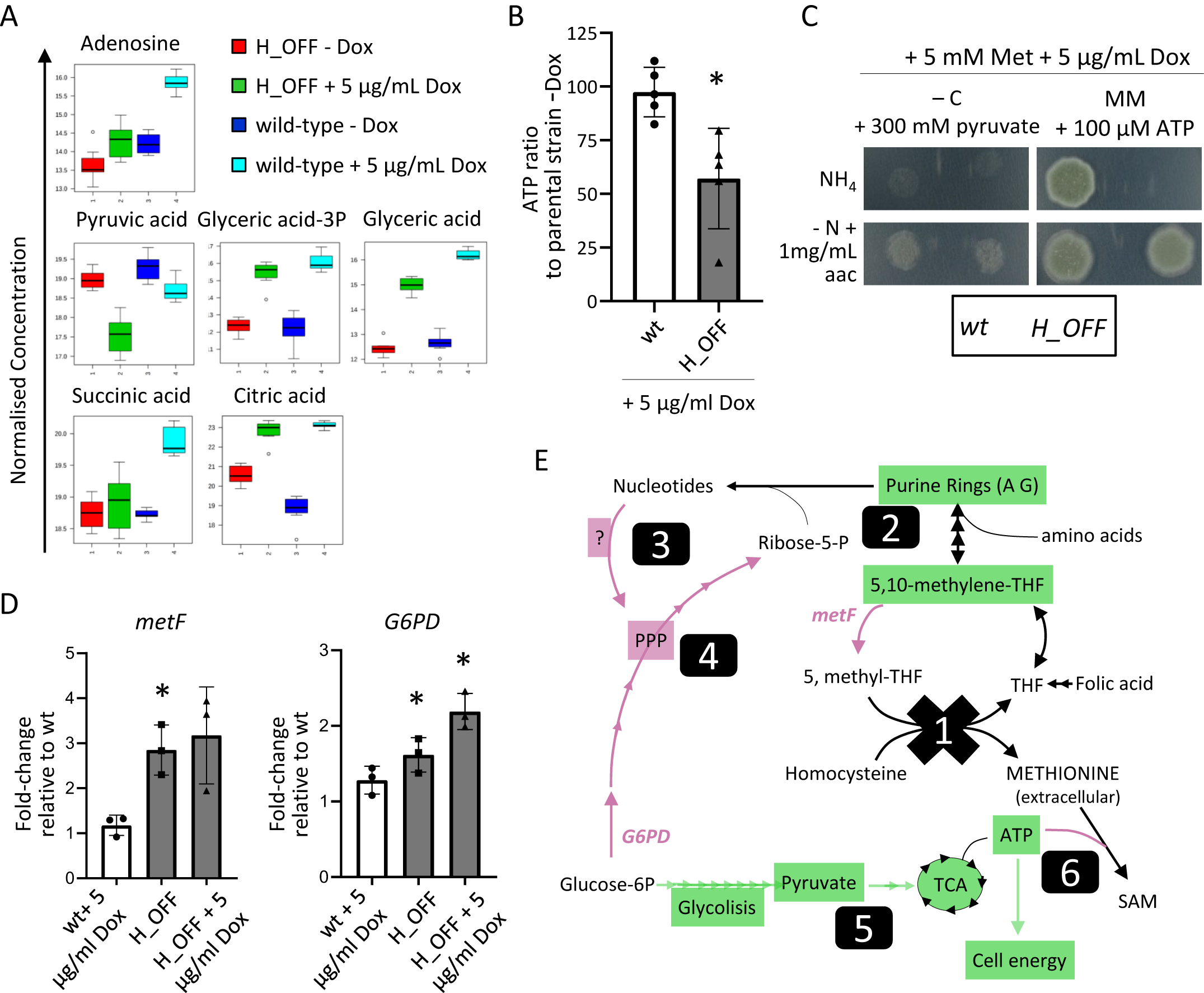
Lack of methionine synthase activity causes a decrease in cell energetics. **A)** Normalized concentrations of metabolites in fungal mycelia (*n*=8). Adenosine levels were decreased in *H_OFF*+Dox compared with *wt*+Dox, which agrees with its capacity to partially reconstitute growth. Several metabolites of the glycolysis and TCA pathways were reduced in *H_OFF*+Dox compared with *wt*+Dox, suggesting low energetic levels. **B)** The levels of ATP significantly decreased in the *H_OFF* strain upon Dox addition, whilst they did not vary significantly in the *wt* strain. Each point represents a biological replicate, which was assayed with three technical replicates. Data was analysed using one sample *t*-test to a hypothetical value of 100 (i.e. no change in ratio of ATP). Graph displays the mean and standard deviation. **C)** Pyruvate as the sole carbon source could reconstitute growth of the *H_OFF* strain in restrictive conditions to wild-type levels. For both strains, growth was limited and slightly improved in the presence of amino acids. ATP could fully reconstitute growth of *H_OFF* when amino acids were the sole N-source. The phenotypic analyses were repeated in three independent experiments. Representative plates are shown. **D)** RT-PCR calculation of the fold-change in genetic expression of *metF* and *G6PD* with respect to their expression in wild-type without Dox. Each point represents a biological replicate which was analysed with three technical replicates. Data was analysed using one sample *t*-test to a hypothetical value of 1 (i.e. no change in expression). Graphs display the mean and standard deviation. **E)** Schematic representation of the metabolic imbalance started with MetH repression (1). Lack of the enzymatic activity caused a shortage of 5,10-methylene-THF and consequently of purine rings (2). This was sensed as a shortage of nucleotides that the cell attempted to compensate through an unknown mechanism, seemingly TOR and PKA independent (3), that activated glucose flow through the Pentose Phosphate Pathway (PPP) (4). That caused a reduction of glycolysis and TCA cycles, which in turn decreased ATP levels (5). ATP usage for *S*-adenosylmethionine (SAM) synthesis was maintained, which caused a drop in cell energy that resulted in growth arrest (6). Genes/pathways/compounds expected to be reduced are highlighted in green and those increased in magenta.

The fact that growth in the absence of methionine synthase can be reconstituted when there are sufficient levels of methionine and ATP implies that *metH* is a conditionally essential gene, meaning that it is only essential in the absence of the specific conditions that overcome the disturbances derived from its deficiency. We believe that a significant number of genes previously described as essential in fungi would in fact be conditionally essential, however the right conditions to reconstitute growth have not been identified in many cases. This highlights a paramount consideration for the proper identification and validation of drug targets: the deficiencies introduced by targeting a conditionally essential gene must not be overcome during infection. This important concept has already been discussed by others (48-51) and we believe addressing it should become the standard for proper validation of antimicrobial targets. In the case of methionine synthase, it is unlikely that the fungus could acquire sufficient levels of ATP (combined with methionine and not using a preferred N-source) in the lung tissue to overcome the growth defect resulting from targeting MetH. The concentration of free extracellular ATP in human plasma has been calculated to be in the sub-micromolar range (28-64 nM) (52). In the lungs, extracellular ATP concentrations must be strictly balanced and increased levels are implicated in the pathophysiology of inflammatory diseases (53); nevertheless, even in such cases ATP levels have been calculated in the low micromolar range (54, 55). Despite this low concentrations and consequently unlikely compensation, we believe that MetH needs to be validated in a suitable model to confirm that its deficiency cannot be not overcome in established infections.

We then questioned how the lack of methionine synthase’s enzymatic activity could cause a drop in cell energy. We hypothesised that blockage of methionine synthase activity likely causes a forced conversion of 5,10-methylene-THF to 5-methyl-THF by the action of MetF (Fig. 1A). In support of this, we observed that expression of *metF* was increased in the *H_OFF* strain (Fig. 3D). This likely causes a shortage of 5,10-methylene-THF, as the conversion is not reversible and THF cannot be recycled by the action of methionine synthase (Fig. 1A). Indeed, supplementation of folic acid (only when amino acids are the sole N-source, Fig. S3) and of purines could partially restore growth (Fig. 2A), as they compensate for the deficit in purine ring biosynthesis when there is a shortage of 5,10-methylene-THF. However, this still does not explain why there is a drop in ATP. We hypothesised that the block of purine biosynthesis might be sensed as a shortage of nucleotides. This could then cause a shift in glucose metabolism from glycolysis and the TCA cycle (which produce energy) to the Pentose Phosphate Pathway (PPP), which is required to produce ribose-5-phosphate, an integral part of nucleotides. In a similar vein, it has recently been described that activation of anabolism in *Saccharomyces cerevisiae* implies increased nucleotide biosynthesis and consequently metabolic flow through the PPP (56). To evaluate our hypothesis, we investigated the transcription level of the glucose-6-phosphate dehydrogenase (G6PD) encoding gene (AFUA_3G08470), which catalyses the first committed step of the PPP. In agreement with our hypothesis, the expression of G6PD encoding gene increases in the *H_OFF* strain upon addition of Dox (Fig. 3D), likely reflecting an increased flow through the PPP. We then wondered how cells may be activating the PPP. The target of rapamycin (TOR) TORC1 effector, which is widely known to activate anabolism and growth (57-59), has been described to activate the PPP in mammalian cells (60, 61) and has been functionally connected with energy production and nucleotide metabolism in *A. fumigatus* (62). In addition, the cAMP/PKA (protein kinase A) pathway is known to be paramount for sensing of nutrients and the correspondent adaptation of gene expression and metabolism (63), and was found to be implicated in the regulation of nucleotide biosynthesis in *A. fumigatus* (64). Consequently, we explored if a partial block of TOR with low concentrations of rapamycin or of PKA with H-89 could prevent the imbalanced activation of the PPP in the absence of MetH activity. However, neither of the inhibitors could reconstitute growth of the *H_OFF* strain in restrictive conditions (Fig. S4C). This means that neither the TOR nor the PKA pathways seem to be involved in the deleterious metabolic shift that seemingly activates PPP and decreases flux through glycolysis. Therefore, more experiments are required to elucidate the mechanism underlying the metabolic imbalance developed upon *metH* downregulation.

In summary, we propose that absence of methionine synthase activity causes a strong defect in purine biosynthesis that the cell tries to compensate for by shifting carbon metabolism to the PPP; this metabolic imbalance causes a drop of ATP levels, which collapses cell energetics and results in halted growth (Fig. 3E).

Interestingly, we also detected that *metF* expression is higher in the *H_OFF* strain compared to the wild-type, even in the absence of Dox (Fig. 3D). This could be explained as an effort to compensate a higher demand of 5-methyl-THF by the slightly increased amount of methionine synthase in this strain (Fig. S1A). This effect could cause a mild defect in purine biosynthesis in the *H_OFF* strain, and indeed adenosine content was lower in the *H_OFF*–Dox condition compared with the wild-type–Dox sample in the metabolome analysis (Fig 3A). Furthermore, this also explains why we detected a small but significant increase of G6PD expression in the *H_OFF*–Dox condition (Fig. 3D). Therefore, it seems that upregulating methionine synthase has the potential to cause the same metabolic imbalance as downregulating it. However, the effect of overexpression (notice that it is only ∼1.5 fold in our strain Fig. S1A) is minor and does not have obvious consequences for growth, as THF can be recycled and thus the shortage of 5,10-methylene-THF is not severe. In any case, two important points can be highlighted from this small imbalance. Firstly, methionine synthase activity is very important and must be finely tuned to maintain a proper metabolic homeostasis. Secondly, changing the expression level of genes with constitutive and/or regulatable promoters can have unexpected and hidden consequences that often go unnoticed.

### Supplementation with S-adenosylmethionine reconstitutes ATP levels and growth

We have shown that the absence of MetH activity causes a reduction in ATP levels. *S*-adenosylmethionine (SAM) is produced from methionine and ATP by the action of *S*-adenosylmethionine synthetase SasA (Fig. 1A), an essential enzyme in *A. nidulans* (65). Hence, we reasoned that the absence of MetH activity might cause a decrease in SAM levels. To test that hypothesis, we first attempted to rescue growth of the *H_OFF* strain in a medium supplemented with methionine and SAM. We tested various N-sources to diversify the presence of permeases in the cell membrane, aiming to maximise the chances of SAM uptake (Fig. S3). Indeed, the addition of SAM reconstituted growth of the *H_OFF* strain in restrictive conditions in the presence of methionine when amino acids were the only N-source (Fig. 4A and S3). We then measured the intracellular concentration of SAM in growing mycelia upon addition of Dox using MS/MS. Surprisingly, we observed that addition of Dox to the *H_OFF* strain did not cause a significant reduction in SAM levels (Fig. 4B). Consequently, we wondered how the addition of SAM may reconstitute *H_OFF* growth if its levels are not reduced upon *metH* repression. We speculated that as SAM is a crucial molecule it continues to be produced even if the levels of ATP are reduced, draining it from other cellular processes and thus triggering energy deprivation. In support of this hypothesis, we observed that supplementing SAM to the medium increased the levels of ATP in growing hyphae (Fig. 4C).

**Figure 4.**
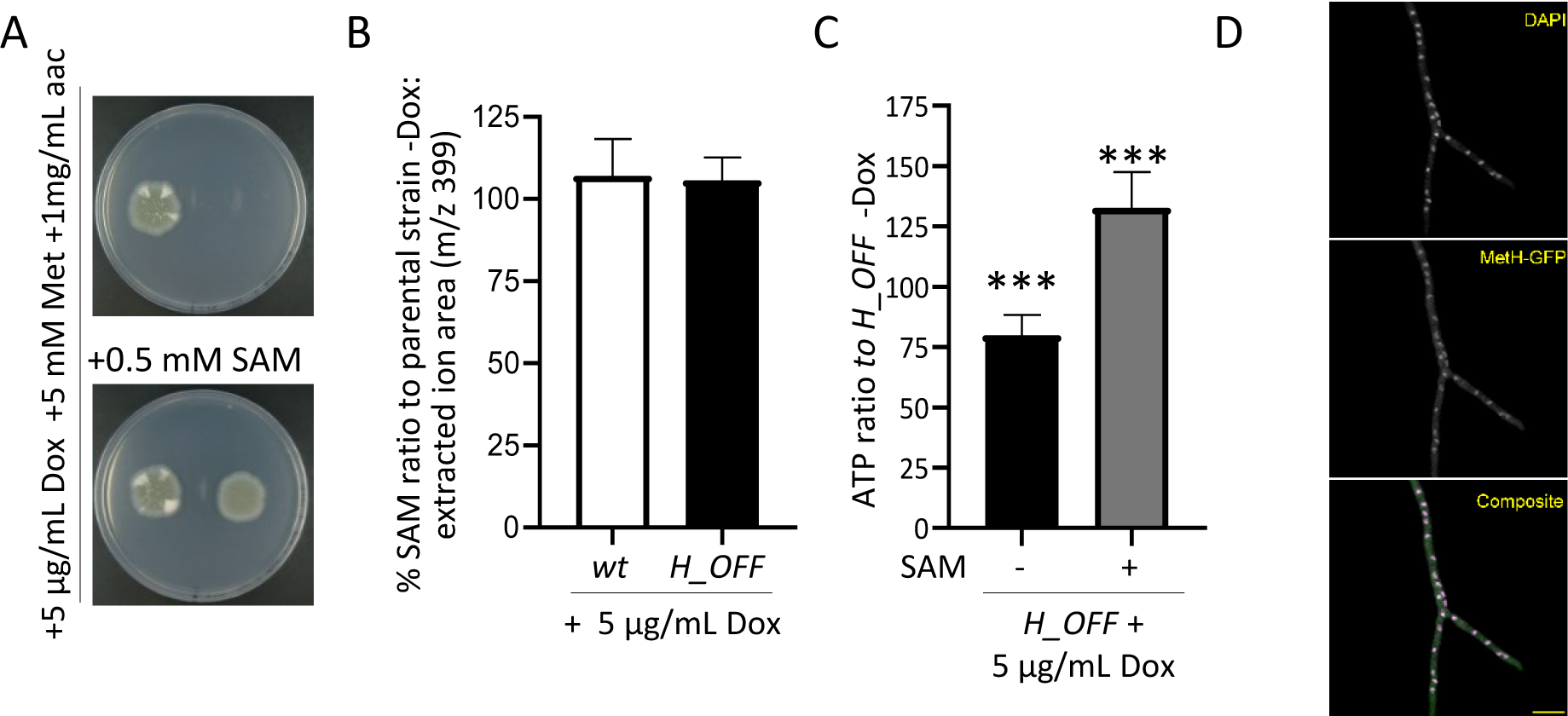
External *S*-adenosylmethionine (SAM) reconstitutes ATP levels and growth. **A)** Addition of SAM to the medium reconstituted growth of the *H_OFF* strain in restrictive conditions. The phenotypic analysis was repeated in two independent experiments. Representative plates are shown. **B)** Levels of SAM did not significantly decrease upon Dox administration (5 µg/mL) in wild-type or *H_OFF* strains. Graphs depict mean and SD of three biological and two technical replicates. Data were analysed using one-way ANOVA with Bonferroni post-tests adjustment. **C)** Presence of SAM in the medium (0.5 mM) prevented the decrease in ATP levels observed in the *H_OFF* strain upon Dox addition. In fact, ATP levels were increased compared to the minus Dox condition. Graphs depict mean and SD of two biological and four technical replicates. **D)** Expression of MetH-GFP in the *H_OFF* background showed that the protein localised both in cytoplasm and nucleus. Singular channels are shown in greyscale. In composite image magenta=DAPI and green=GFP (Bar=10 µm).

As SAM supplementation can reconstitute *H_OFF* growth, it constitutes another condition that overcomes the conditional essentiality of *metH*, highlighting again the need to validate MetH in a model of established infection. The concentration of SAM in human serum is extremely low, in the range of 100-150 nM (30), and consequently it is unlikely that the fungus could find sufficient SAM during infection to compensate for the defect in ATP caused by targeting methionine synthase.

*S*-adenosylmethionine plays a fundamental role as methyl donor for the majority of cellular methylation reactions, including methylation of DNA. Given the observed importance of SAM in the absence of MetH activity and considering that in *P. pastoris* and *C. albicans* methionine synthase was reported to localise in the nucleus, as well as in the cytoplasm (33), we speculated that nuclear localization might be important for MetH cellular function. To test this hypothesis, we constructed strains expressing different versions of C-terminus GFP-tagged MetH from the pJA49 plasmid (Fig. S2) in the *H_OFF* background. These were a wild-type MetH, a MetH^D616A^ (control of no growth –Fig. 1C & S5A–) and a MetH^R749A^ (*metH*^*g2439CG>GA*^) version of the protein, which according to the results published for *P. pastoris* should not localise in the nucleus (33). The strain expressing wild-type MetH grew normally in restrictive conditions (Fig. S5A), proving that the tagged MetH-GFP protein was active. Importantly, this result also demonstrated that genetic downregulation of *metH* is the only reason for the lack of growth of the *H_OFF* strain in the presence of Dox. We confirmed that *A. fumigatus* MetH localises in both the nucleus and cytoplasm (Fig 4D & S5B). In contrast to what was described in *P. pastoris*, the MetH^R749A^ protein seems to be active, as it could trigger growth of *H_OFF* in restrictive conditions (Fig. S5A) and still localised into the nucleus (Fig. S5B). Therefore, the possibility that MetH localisation in the nucleus is important needs further exploration.

### Repression of methionine synthase causes growth inhibition in growing mycelia

The major advantage of the tetOFF system is that it can be employed to simulate a drug treatment before a specific chemical is developed. Addition of Dox to a growing mycelium downregulates the gene of interest (Fig S1A), mimicking the effect of blocking its product by the action of a drug. The validity of the Tet systems has recently been questioned, as it has been reported that Dox can impair mitochondrial function in various eukaryotic models (66). Nevertheless, existing evidence suggests that low concentrations of Dox (≤ 50 µg/mL) have little effect on fungal cells. For instance, in contrast to reports of Dox affecting proliferation of human cells at low concentrations (66, 67), we and others have not detected negative effects of low concentrations of Dox or of tetracycline on fungal proliferation (Fig. 5A), phenotype (macroscopic or microscopic, Fig. 5B) or virulence (17, 68-74). In addition, it was reported that 40 µg/mL of Dox does not affect the transcriptional profile of *S. cerevisiae* (75), which contrasts with the broad effect caused by only 1 μg/mL Dox on global transcription in human cells (66). Moreover, a study that investigated the role of various mitochondrial proteins for its function in *C. albicans* did not observe any negative effect of 20 µg/mL Dox on fungal growth nor on mitochondrial morphology and function (76). To test the effect of Dox on *A. fumigatus*, we grew the wild-type strain overnight in various concentrations of drug and imaged mitochondria using the Rhodamine 123 dye (Fig S6). It was previously described that inhibition of translation in mitochondrial (mechanism of Dox toxicity) promotes mitochondrial fission, which can be detected as a more fragmented, punctuate, mitochondrial appearance compared to the healthy tubular morphology (66, 76). This fragmented phenotype started to appear, although there was variation among hyphae, when the fungus was incubated in 100 µg/mL Dox and became obvious when it was incubated in 1000 µg/mL Dox (Fig. S6). In contrast, in low concentrations of Dox (1 and 10 µg/mL Dox) the mitochondria showed a healthy tubular morphology, indistinguishable from that of no Dox (Fig. S6). Therefore, low concentrations of Dox do not affect mitochondria morphology and thus likely do not impair their function. In fact, we have also observed that addition of 5 µg/mL Dox to wild-type mycelium did not affect the ATP content (Fig 3B), further supporting the conclusion that mitochondrial function is not impaired. Therefore, even if higher concentrations of Dox have a negative effect on fungal cells (70, 74), its impact at low concentrations on fungal cells seems to be minimal. Consequently, we argue that as long as the concentration of Dox used is ≤ 50 µg/mL, the tetOFF system can be utilised to investigate the consequences of downregulating gene expression in fungal research.

**Figure 5.**
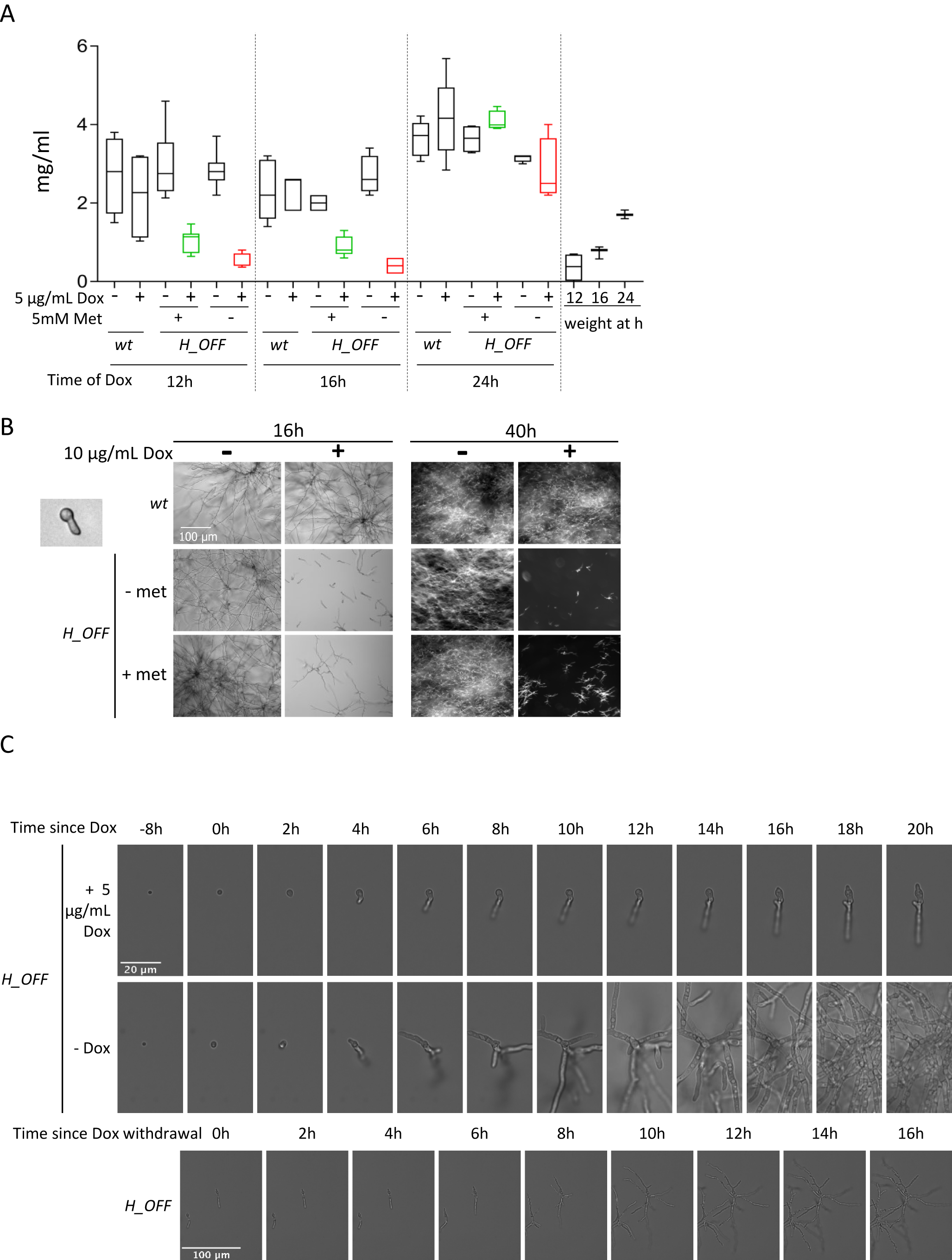
Repression of *metH* transcription causes inhibition of growth *in vitro*. **A)** Addition of 1 µg/mL Dox to 12 and 16 h grown mycelia strongly reduced fungal biomass, measured 24 h later. This was more pronounced in media without methionine. The effect was lost when Dox was added 24 h after inoculation. For comparison, fungal biomass of mycelia harvested at the time of Dox addition (12, 16 and 24 h post-inoculation) is shown. Three independent experiments were performed, using 3 technical replicates for each. **B)** Microscopic images of 8 h germinated spores treated with 10 µg/mL Dox. Images were taken 16 h after Dox addition (wide-field microscopy) and again 24 h later (stereomicroscopy). Dox addition halts growth of *H_OFF* strain in a sustained manner. **C)** Time-lapse microscopy of *H_OFF* growth upon Dox addition. Dox was added to 8 h grown conidia, which caused growth inhibition that was obvious after ∼4 h. Growth was virtually halted for as long as Dox was present. **D)** ∼6 h after Dox withdrawal, *H_OFF* growth resumed, showing that the effect was fungistatic.

To investigate the effect of downregulating *metH* for mycelial growth, we added Dox to 12, 16 or 24 h grown submerged mycelia and left it incubating for an additional 24 h. Addition of Dox to 12 or 16 h grown mycelia severely impaired growth of the *H_OFF* strain but not the wt, as observed by biomass (Fig. 5A) and OD (Fig. S7A) measurements. This effect was lost when Dox was added to 24 h grown mycelia, due to the incapacity of Dox to reach and downregulate expression in all cells within the dense mass of an overgrown mycelium. Interestingly, Dox addition to methionine free media stopped *H_OFF* growth immediately, which can be observed by comparing fungal biomass at the time of Dox addition to the measurement 24 h after Dox addition. In contrast, the fungus inoculated in methionine containing media grew a little further after Dox addition (Fig. 5A). To understand this difference, we added Dox to either resting or 8 h germinated conidia and imaged them 16 and 40 h after drug addition (Fig. 5B & Fig. S7B). In agreement with the previous result, we observed that Dox addition in methionine free medium inhibited growth immediately: resting conidia did not germinate and germinated conidia did not elongate the germtube. In contrast, after addition of Dox in methionine containing medium, most of the resting conidia were still able to germinate and some germlings could elongate the germinated tubes to form short hyphae. This suggests that the drop in ATP levels takes ∼3-4 h before having an effect on growth. Importantly, once growth was inhibited, the effect was sustained for a long period, as we could not detect further growth up to 40 h post-inoculation. To corroborate these observations and further determine whether the effect of growth is fungistatic or fungicidal in the long term, we performed a time-lapse analysis of the effects of adding Dox to 8 h swollen conidia and its subsequent withdrawal after 16 h of incubation (Fig. 5C and Video 1). We observed that growth was inhibited ∼4 h after Dox addition and almost completely halted after 6 h, which was sustained as long as the drug was present. Upon withdrawal of Dox, growth resumes within 6 h (Fig 5D and Video S1), showing that the effect of blocking MetH is fungistatic, at least with the genetic TeOFF model of *metH* repression. As expected Dox had no effect on wild-type growth (Video S2).

### Targeting MetH in established infections interferes with the progression of disease

We previously used the TetON system to investigate the relevance of MetH in *A. fumigatus* virulence (17). In this model, *metH* gene expression was active when mice were fed with Dox, which resulted in full virulence of the *metH_tetON* strain (demonstrating that the concentration of Dox reached in murine tissues does not impact fungal virulence). In the absence of Dox the gene was not expressed, which completely abrogated virulence. This proved that the murine lung does not readily provide the conditions to overcome the conditional essentiality of *metH*, and thus this gene is required to establish infection. However, antifungal drugs are normally administered to treat patients who already have an established infection. Therefore, it is possible that the conditional essentiality of the gene could be overcome when the fungus is actively growing in the tissue, as the fungal metabolic requirements and the environmental conditions are different (49, 50). Consequently, in order to achieve a rigorous target validation, it is crucial to assess the efficiency of new target candidates in established infections. The first conditional promoter system used for *A. fumigatus in vivo* was the (p)niiA (48). This system represses genetic expression of the gene of interest in the presence of ammonium, which is contained in murine serum. Accordingly, in this seminal study several genes essential *in vitro* were confirmed or refuted to be essential to initiate infection in a model of systemic (blood) infection. Currently, two more systems are in use to assess the relevance of fungal essential genes for pulmonary infection: TetON and (p)xylP. These systems can be used to either impede or permit fungal gene expression in murine lungs; but in both models this control can only be exerted from the beginning of infection, as sufficient levels of the inducing molecule (doxycycline or xylose) must be present to activate gene expression in the control condition. Consequently, these models have been used to investigate the relevance of genes required to grow *in vitro* to initiate pulmonary infection (17, 72, 77). However, those models cannot be used to determine the importance of the genes in established aspergillosis infections *in vivo*. Therefore, we aimed to optimise the use of the TetOFF system for this purpose, as it can be used to downregulate gene expression in growing mycelia. As a control for the model, we constructed a *cyp51A_tetOFFΔcyp51B* (*51A_OFF*) strain. We reasoned that the target of the azoles, first-line treatment drugs for *Aspergillus* diseases, should be the gold standard to compare any target against. This strain showed a similar behaviour as *H_OFF in vitro*: as little as 0.05 µg/ml Dox prevented colony development on an agar plate (Fig. S8A) and addition of Dox to conidia or germlings blocked growth (Fig. S8B).

We first assayed the use of the TetOFF system in the *Galleria mellonella* alternative mini-host model of infection. Preliminary experiments revealed that the balance between reaching sufficient levels of Dox to exert an effect and maintaining toxic effects of overdose low was very delicate. We finally optimised a regimen consisting of 5 injections of 50 mg/kg Dox (Fig. S9A) that caused little mortality in the control group (25% in Dox control VS 12% in PBS treatment control *P*=0.22) but still showed an effect of treatment (Fig. 6A). We then infected *Galleria* larvae with 5×10^2^ conidia of *51A_OFF* or *H_OFF* strains and applied the Dox regimen or PBS vehicle starting at the same time of infection (0 h) or 6 h after infection (Fig. S9A). For both strains, administration of Dox from the beginning of infection triggered a significant improvement in survival compared with the non-treated conditions (50% VS 17.2% for *51A_OFF, P*=0.0036, and 41.45% VS 6.67% for *H_OFF, P*=0.022) (Fig. 6A). The fact that administration of Dox at the time of infection did not improve survival to close to 100%, was not surprising, as it is important to note that Dox does not completely prevent gene expression (Fig. S1A), so a moderate effect on survival was expectable. Furthermore, rapid metabolization of the drug in the larvae hemocoel or microenvironment variations in its concentration may also account for a discrete effect of treatment. Despite this limitations of the model, we observed that administration of Dox 6 h after infection also triggered a significant improvement in survival for both strains (42.8% VS 17.2% for *51A_OFF, P*=0.0007, and 32.26% VS 6.67% for *H_OFF, P*=0.0324) (Fig. 6A). Therefore, downregulation of methionine synthase genetic expression in established infections conferred a significant benefit in survival which was comparable to that observed with the target of the azoles.

**Figure 6.**
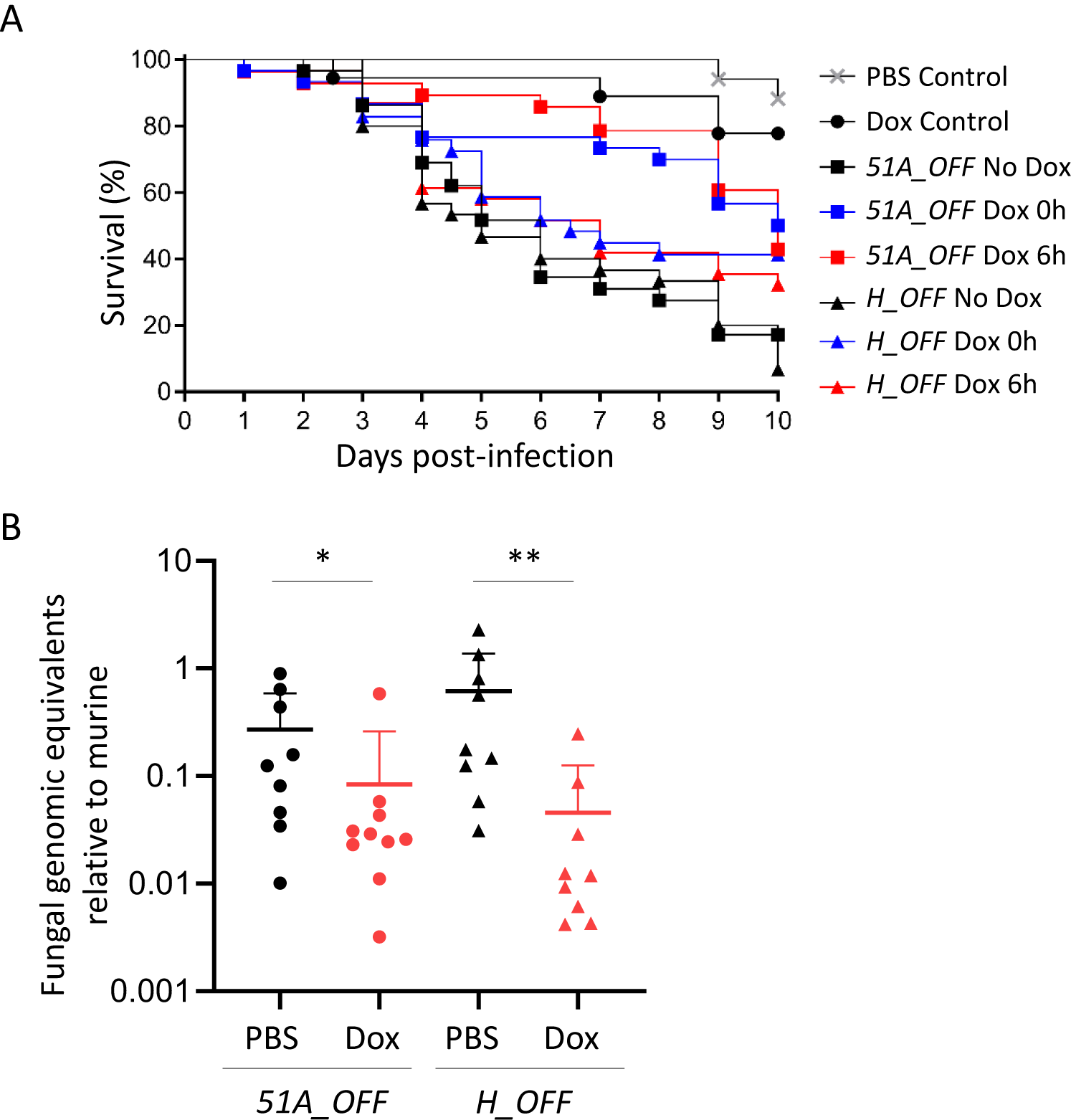
Downregulation of *metH in vivo* in established infections shows a beneficial effect comparable to the target of azoles. **A)** Administration of a Dox regimen (Fig. S9A) to *Galleria mellonella* infected with either the *H_OFF* strain or the *51A_OFF* control strain showed a beneficial effect in survival. For both strains, starting regimen at the time of infection triggered a significant improvement in survival (50% VS 17.2% for *51A_OFF, P*=0.0036, and 41.45% VS 6.67% for *H_OFF, P*=0.022). Dox regimen 6 h after infection also triggered a significant improvement in survival for both strains (42.8% VS 17.2% for *51A_OFF, P*=0.0007, and 32.26% VS 6.67% for *H_OFF, P*=0.0324). The curves show the pooled data from 3 independent experiments. Curves were compared using the Log-Rank test. **B)** Leukopenic mice were infected with *51A_OFF* or *H_OFF* and a Dox regimen (Fig. S9B) administered 16 h after infection. Upon Dox treatment, fungal burden in the lungs was significantly reduced for both strains (*P*=0.0279 for *51A_OFF* and *P*=0.0019 for *H_OFF*). Two independent experiment were carried out. Each point in the graphs represents one mouse (n=9). Burdens for each strain were compared using a Mann Whitney test.

The positive results obtained using the *Galleria* infection model prompted us to assay the TetOFF system in a leukopenic murine model of pulmonary aspergillosis. To ensure that Dox levels in mouse lungs reach and maintain sufficient concentrations to downregulate gene expression (according to our results *in vitro*) we performed a pilot Dox dosage experiment in immunosuppressed non-infected mice (Fig S8B). We extracted lungs of Dox treated mice at different time-points, homogenated them and measured Dox concentration using a bioassay based on inhibition of *Escherichia coli DH5α* growth. We could detect promising levels of Dox in all mice (concentrations ranging from 2.2 to 0.94 µg/mL –Fig. S9B–) which according to our results *in vitro* should be sufficient to downregulate gene expression from the TetOFF system. We therefore infected leukopenic mice with 10^5^ spores of the *51A_OFF* or the *H_OFF* strains and administered PBS vehicle or our Dox regimen, starting 16 h after infection (Fig. S9B). The use of an uninfected, Dox treated control group uncovered that the intense Dox regimen used was harmful for the mice. These uninfected mice lost weight at a similar rate as the infected groups and looked ill from the third or fourth day of treatment. This is not surprising as Dox can impair mitochondrial function in mice (66) and has iron chelating properties (78). As a consequence, there was no beneficial effect of Dox treatment on survival (not shown). The fact that Dox treatment did also not show any benefit in survival for our control strain *51A_OFF*, which should mimic treatment with azoles (primary therapy for invasive aspergillosis), indicates that the TetOFF system is not ideal to mimic a drug treatment in established infections. Nevertheless, we further attempted to determine the efficiency of targeting MetH in established infections by measuring fungal burdens in lungs of treated and untreated mice. We observed that two and a half days of Dox treatment (when the mice have not developed visible toxic effects yet) did result in a significant reduction of fungal burdens 3 days after infection for both *51A_OFF* (*P*=0.0279) and *H_OFF* (*P*=0.0019) (Fig. 6B). Therefore, we could observe a beneficial effect of interfering with methionine synthase genetic expression in an established pulmonary infection, which was comparable to that of interfering with the expression of *cyp51A*, the target of azoles. This constitutes a very rigorous validation of MetH as a promising antifungal target.

A recent study also aimed to use another TetOFF system to validate a drug target in established aspergillosis infections (79). These authors administered Dox exclusively through oral gavage, accounting for lower dosage of drug. Consequently, even if no toxic effect for the mice was observed, they also did not detect any beneficial effect on survival when the Dox treatment was initiated after infection. Therefore, the TetOFF system is clearly not optimal and better models are needed. Yet, it is currently the only model with which the efficiency of new targets can be tested in aspergillosis established infections, and thus it is highly valuable that we have been able to optimise its use *in vivo*. Our *51A_OFF* control strain has been key to calibrating the model, and allows us to be confident that the beneficial effects observed, even if subtle, are significant. Hence, we propose that hereinafter proper genetic validation of antifungal targets should include testing their relevance in established infections.

### Structural-based virtual screening of MetH

Having shown *in vivo* that MetH is a promising target, we decided to investigate its druggability by running a structural-based virtual screening. The sequence of *A. fumigatus* MetH (AfMetH) contains two predicted methionine synthase domains with a β-barrel fold conserved in other fungal and bacterial enzymes. The structure of the *C. albicans* orthologue (80) (CaMetH) showed that the active site is located between the two domains where the methyl tetrahydrofolate, the homocysteine substrate and the catalytic zinc ion bind in close proximity. The homology model for AfMetH (Fig. 7A) overlaps very well with that of the CaMetH thus providing a suitable molecular model for further analysis. In contrast, the structure of the human methionine synthase (hMS) shows a very different overall arrangement with the folate and homocysteine binding domains located in completely different regions (Fig. 7B). Comparison of the tetrahydrofolate binding sites between the fungal and the human structures also highlights significant structural differences that affect the conformation adopted by the ligand. In the CaMetH structure the 5-methyl-tetrahydrofolate (C2F) adopts a bent conformation (<20Å long) and it is in close proximity to the methionine product, whereas in the human structure the tetrahydrofolate (THF) ligand binds in an elongated conformation extending up to 30Å from end to end, (Fig. 7 C&D).

**Figure 7.**
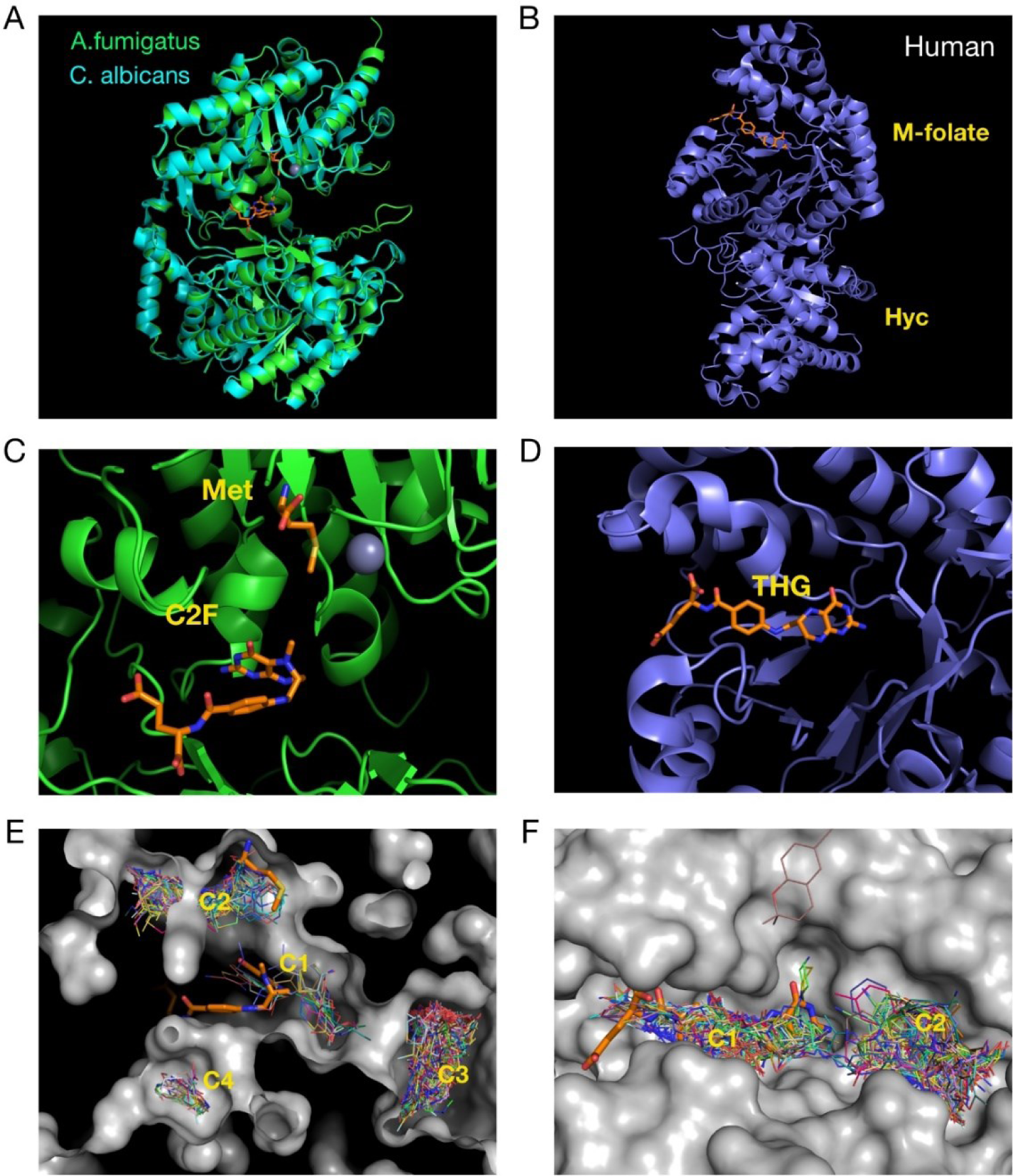
Virtual-screening of fungal and human methionine synthases reveals different druggability of the proteins. **A)** The structures of the crystallized *C. albicans* and the predicted A. *fumigatus* methionine synthases are highly similar. **B)** In contrast, the structure of the human enzyme is very different, having he 5-methyl-tetrahydrafolate and homocysteine binding sites separated. **C)** Detail of the active site of fungal methionine synthases, with the methionine, folate and zinc displayed. **D)** Tetrahydrofolate binding site of the human protein. **E)** Virtual screening on the *A. fumigatus* protein found four ligand binding clusters in the structure, two of which (C1, C2) match the binding position of the 5-methyl-tetrahydrafolate and the methionine. **F)** In the human enzyme two clusters were found, one (C1) that overlaps with the tetrahydrofolate binding site and (another C2) in a nearby pocket.

Virtual screening (VS) was carried on the AfMetH and the hMS structures with the Maybridge Ro3 fragment library to explore potential venues for drug development. The results showed four ligand binding clusters in the AfMetH structure, two of which (C1, C2) match the binding position of the 5-methyl-tetrahydrafolate and the methionine from the CaMetH crystal structure (Fig. 7E). For the hMS, we found two main clusters, C1 that overlaps with the tetrahydrofolate binding site and C2 in a nearby pocket. Clearly the distribution of the clusters defines a very different landscape around the folate site between the human and the fungal enzymes. Furthermore, the proximity of the C1 and C2 clusters, matching the folate and Met/homocysteine binding sites in the Ca/Af proteins means that it may be possible to combine ligands at both sites to generate double-site inhibitors with high specificity towards the fungal enzymes. Antifolates are a class of drugs that antagonise folate, blocking the action of folate dependent enzymes such as dihydrofolate reductase (DHFR), thymidylate synthase or methionine synthase. Methotrexate is an antifolate commonly used to treat cancer and autoimmune diseases. Interestingly, methotrexate has been shown to be a weak inhibitor of the *C. albicans* methionine synthase (32) and to have some antifungal activity against *C. albicans* (81) and *Aspergillus ssp* (82). Nevertheless, methotrexate is not a good antifungal drug, as its activity is high against human enzymes (IC_50_ of 0.3 µM for DHFR (83)) and low against fungal methionine synthase (IC_50_ of 4 mM for *C. albicans* MetH (32)). Therefore, more potent and specific inhibitors of fungal methionine synthases are needed to fully exploit the value of this target for antifungal therapy, a task that seems possible and can be directed from our analyses.

In summary, we have shown that methionine synthase blockage triggers not only methionine auxotrophy, but also a metabolic imbalance that results in a drop in cellular energetics and growth arrest. In light of our results, we stress that conditional essentiality is important to understand the underlying mechanisms of metabolic processes and needs to be considered to achieve proper validation of novel antimicrobial targets. Accordingly, we proved that targeting methionine synthase in established infections has a beneficial effect similar to that observed for the target of azoles, the most effective drugs for the treatment of aspergillosis. Finally, we showed that fungal methionine synthases have distinct druggable pockets that can be exploited to design specific inhibitors. In conclusion, we have demonstrated that fungal methionine synthases are promising targets for the development of novel antifungals.

## MATERIAL AND METHODS

### Strains, media and culture conditions

The *Escherichia coli* strain DH5α (84) was used for cloning procedures. Plasmid-carrying *E. coli* strains were routinely grown at 37°C in LB liquid medium (Oxoid) under selective conditions (100 µg·mL^-1^ ampicillin or 50 µg·mL^-1^ kanamycin); for growth on plates, 1.5% agar was added to solidify the medium. All plasmids used in the course of this study were generated using the Seamless Cloning (Invitrogen) technology as previously described (17, 85). *E. coli* strain BL21 (DE3) (86) was grown on Mueller Hinton agar (Sigma) in bioassays, to determine Dox concentrations within homogenized murine lungs.

The wild-type clinical isolate *Aspergillus fumigatus* strain ATCC 46645 served as reference recipient. *A. fumigatus* strain A1160 (*ku80Δ*) (87) was also used to confirm *metH* essentiality. *A. fumigatus* mutants were generated using a standard protoplasting protocol (88). *A. fumigatus* strains were generally cultured in minimal medium (MM) (89) (1% glucose, 5 mM ammonium tartrate, 7 mM KCl, 11 mM KH_2_PO_4_, 0.25 mM MgSO_4_, 1× Hutner’s trace elements solution; pH 5.5; 1.5% agar) at 37°C. For selection in the presence of resistance markers 50 µg·mL^-1^ of hygromycin B or 100 µg·mL^-1^ of pyrithiamine (InvivoGen) were applied. In sulfur-free medium (MM-S), MgCl_2_ substituted for MgSO_4_ and a modified mixture of trace elements lacking any sulfate salt was used. For all growth assays on solid media, the culture medium was inoculated with 10 µl of a freshly prepared *A. fumigatus* spore suspension (10^5^ conidia·mL^-1^ in water supplemented with 0.9% NaCl and 0.02% Tween 80) and incubated at 37°C for 3 days.

### Extraction and manipulation of nucleic acids

Standard protocols of recombinant DNA technology were carried out (90). Phusion^®^ high-fidelity DNA polymerase (ThermoFisher Scientific) was generally used in polymerase chain reactions and essential cloning steps were verified by sequencing. Fungal genomic DNA was prepared following the protocol of Kolar *et al*. (91) and Southern analyses were carried out as described (92, 93), using the Amersham ECL Direct Labeling and Detection System^®^ (GE Healthcare). Fungal RNA was isolated using TRIzol reagent (ThermoFisher Scientific) and Qiagen plant RNA extraction kit. Retrotranscription was performed using SuperScript III First-Strand Synthesis (ThermoFisher Scientific). RT-PCR on both gDNA and cDNA was performed using the SYBR® Green JumpStart (Sigma) in a 7500 Fast Real Time PCR cycler from Applied Biosystems.

### Microscopy

10^3^ *A. fumigatus* resting or 8 h germinated conidia were inoculated in 200 µL of medium (+/-Dox) in 8 well imaging chambers (ibidi) and incubated at 37°C. Microscopy images were taken on a Nikon Eclipse TE2000-E, using a CFI Plan Apochromat Lambda 20X/0.75 objective and captured with a Hamamatsu Orca-ER CCD camera (Hamamatsu Photonics) and manipulated using NIS-Elements 4.0 (Nikon). For extensively grown mycelia a stereomicroscope Leica MZFL-III was used, with a Q-imaging Retinga 6000 camera, and manipulated using Metamorph v7760. Confocal imaging was performed using a Leica TCS SP8x inverted confocal microscope equipped with a 40X/0.85 objective. Nuclei were stained with DAPI (Life Technologies Ltd) as described previously (94). GFP was excited at 458 nm with an Argon laser at 20% power. DAPI was excited at 405 nm with an LED diode at 20%.

### Metabolome analyses

*A. fumigatus* wild-type and *metH_tetOFF* strains were incubated in MM for 16 h before the −Dox samples were taken (8 replicates of 11 mL each). Then, 5 µg/mL Dox and 5 mM methionine (to prevent metabolic adaptation due to met auxotrophy) were added as appropriate and the cultures incubated for 6 h, after which the +Dox samples were taken (8 × 11 mL). The samples were immediately quenched with 2× volumes of 60% methanol at −48°C. After centrifugation at 4800 *g* for 10 min at −8°C, metabolites were extracted in 1 mL 80% methanol at −48°C by three cycles of N_2_ liquid snap freezing, thawing and vortexing. Supernatant was cleared by centrifugation at −9 °C, 14,500 *g* for 5 min. Quality control (QC) samples were prepared by combining 100 µL from each sample. Samples were aliquoted (300 µL), followed by the addition of 100 µL of the internal standard solution (0.2 mg/mL succinic-*d*_4_ acid, and 0.2 mg/mL glycine-*d*5) and vortex mix for 15 s. All samples were lyophilised by speed vacuum concentration at room temperature overnight (HETO VR MAXI vacuum centrifuge attached to a Thermo Svart RVT 4104 refrigerated vapour trap; Thermo Life Sciences, Basingstoke, U.K.). A two-step derivatization protocol of methoxyamination followed by trimethylsilylation was employed (95).

GC-MS analysis was conducted on a 7890B GC coupled to a 5975 series MSD quadrupole mass spectrometer and equipped with a 7693 autosampler (Agilent, Technologies, UK). The sample (1 μL) was injected onto a VF5-MS column (30 m × 0.25 mm × 0.25 μm; Agilent Technologies) with an inlet temperature of 280 °C and a split ratio of 20:1. Helium was used as the carrier gas with a flow rate of 1 mL/min. The chromatography was programmed to begin at 70 °C with a hold time of 4 min, followed by an increase to 300 °C at a rate of 14 °C/min and a final hold time of 4 min before returning to 70 °C. The total run time for the analysis was 24.43 min. The MS was equipped with an electron impact ion source using 70 eV ionisation and a fixed emission of 35 μA. The mass spectrum was collected for the range 50-550 m/z with a scan speed of 3,125 (*N*=1). Samples were analysed in a randomised order with the injection of a pooled biological quality control sample after every 6th sample injection.

For data analysis, the GC-MS raw files were converted to mzXML and subsequently imported to R. The R package “erah” was employed to de-convolve the GC-MS files. Chromatographic peaks and mass spectra were cross-referenced with the Golm library for putative identification purposes, and followed the metabolomics standards initiative (MSI) guidelines for metabolite identification (96). The peak intensities were normalised according to the IS (succinic-*d*_*4*_ acid) before being log_10_-scaled for further statistical analysis. All pre-processed data were investigated by employing principal component analysis (PCA) (97).

The raw data of this metabolome analysis has been deposited in the MetaboLights database (98), under the reference MTBLS1636 (www.ebi.ac.uk/metabolights/MTBLS1636)

### ATP Quantitation

*A. fumigatus* was grown as in the metabolome analysis. However, where the effect of SAM was investigated spores were inoculated into MM-N + 1mg/mL aac and 0.5mM SAM was also added at the time of Dox addition. ATP levels were determined using the BacTiter-Glo™ Assay (Promega) following the manufacturer’s instructions and a TriStar LB 941 Microplate Reader (Berthold).

### Isolation and detection of SAM

*A. fumigatus* was grown exactly in the same conditions as described for the metabolome analysis. Harvested mycelia were snap-frozen in liquid N_2_ and stored at −70 °C before SAM isolation. SAM extraction was carried out according to Owens *et al* (99). Briefly, frozen mycelia were ground in liquid N_2_ and 0.1 M HCl (250 µL) was added to ground mycelia (100 mg). Samples were stored on ice for 1 h, with sample vortexing at regular intervals. Samples were centrifuged at 13,000 *g* for 10 min (4 °C) to remove cell debris and supernatants were collected. Concentration of protein in supernatants was determined using a Biorad Bradford protein assay relative to a bovine serum albumin (BSA) standard curve. Clarified supernatants were adjusted to 15 % (w/v) trichloroacetic acid to remove protein. After 20 min incubation on ice, centrifugation was repeated and clarified supernatants were diluted with 0.1 % (v/v) formic acid. Samples were injected onto a Hypersil Gold aQ C18 column with polar endcapping on a Dionex UltiMate 3000 nanoRSLC with a Thermo Q-Exactive mass spectrometer. Samples were loaded in 100 % Solvent A (0.1 % (v/v) formic acid in water) followed by a gradient to 20 % B (Solvent B: 0.1 % (v/v) formic acid in acetonitrile) over 4 min. Resolution set to 70000 for MS, with MS/MS scans collected using a Top3 method. SAM standard (Sigma) was used to determine retention time and to confirm MS/MS fragmentation pattern for identification. Extracted ion chromatograms were generated at m/z 399-400 and the peak area of SAM was measured. Measurements were taken from three biological and two technical replicates per sample, normalized to the protein concentration in the extracts from each replicate. SAM levels are expressed as a percentage relative to the parental strain in the absence of Dox.

### Nuclei isolation and Western Blot

Protoplasts were generated as in *A. fumigatus* transformations and nuclei isolated were isolated by sucrose gradient fractionation as previously described by Sperling and Grunstein (100). Nuclear localisation of GFP-tagged target proteins was confirmed by Western-blot. Aliquots of nuclei were boiled for 5 minutes in loading buffer (0.2 M Tris-HCl, 0.4 M DTT, 8% SDS, trace bromophenol blue) and separated on a 12% (w/v) SDS-PAGE gel. The proteins were transferred to a Polyvinylidene difluoride (PVDF) membrane using the Trans-Blot® Turbo™ Transfer System (Bio-Rad). Detection of GFP was carried out with a rabbit polyclonal anti-GFP antiserum (Bio-Rad) and anti-rabbit IgG HRP-linked antibody (Cell Signalling Technology). SuperSignal West Pico PLUS Chemiluminescent Substrate (Thermo Scientific) and the ChemiDoc XRS+ Imaging System (Biorad) were used to visualise immunoreactive bands. Ponceau S staining was performed to normalize the Western-blot signal to the protein loading.

### Mitochondria imaging

Approximately 200 *A. fumigatus* spores were seeded onto Ibidi 8-well slides in minimal medium containing 0, 1, 10, 100 and 1000 µg/ml Dox and incubated at 37 °C for 16 hours. The culture medium was then replaced with minimal medium containing the same Dox concentrations plus 10 µM Rhodamine 123 dye and further incubated for 1-2 hours at 37 °C prior to live-cell confocal imaging. High-resolution confocal fluorescence imaging was performed at 37 °C on a Leica SP8x using a 63x/1.3NA oil immersion lens, whereby Rhodamine 123 fluorescence was excited with a white light laser (5%) tuned to 508 nm and the emission collected on HyD detectors set to 513-600 nm. Representative single plane images of each condition were background-subtracted (rolling ball 20 pixel radius) in Fiji (101) and montaged using FigureJ (102).

### Biomass measurement

Conidia were inoculated into MM-S, supplemented with either methionine or sulfate, and incubated at 37°C 180 rpm for 12, 16 or 24 h. After this initial incubation, 3 mL samples were taken in triplicate from the cultures, filtered through tared Miracloth, dried at 60°C for 16 h and their biomass measured. In treated conditions Dox was added to a final concentration of 1 μg/mL and the culture allowed to grow for a further 24 h at 37°C 180 rpm. 5 mL samples were taken in triplicate and their biomass measured as above.

### Galleria mellonella infections

Sixth-stage instar larval *G. mellonella* moths (15 to 25 mm in length) were ordered from the Live Foods Company (Sheffield, United Kingdom). Infections were performed according to Kavanagh and Fallon (103). Randomly selected groups of 15 larvae were injected in the last left proleg with 10 µL of a suspension of 5×10^4^ conidia/mL in PBS, using Braun Omnican 50-U 100 0.5-mL insulin syringes with integrated needles. Dox was administered according to the treatment shown in Fig. S9A, alternating injections in the last right and left prolegs. In each experiment an untouched and a saline injected control were included, to verify that mortality was not due to the health status of the larvae or the injection method. Three independent experiments were carried out. The presented survival curves display the pooled data, which was analysed with the Log-Rank test.

### Ethics Statement

All mouse experiments were performed under United Kingdom Home Office project license PDF8402B7 and approved by the University of Manchester Ethics Committee and by the Biological Services Facility at the Faculty of Biology, Medicine and Health, University of Manchester.

### Leukopenic murine model of invasive pulmonary aspergillosis and calculation of fungal burden

Outbred CD1 male mice (22– 26 g) were purchased from Charles Rivers and left to rest for at least 1 week before the experiment. Mice were allowed access *ad libitum* to water and food throughout the experiment. Mice were immunosuppressed with 150 mg/kg of cyclophosphamide on days −3 and −1 and with 250 mg/kg cortisone acetate on day −1. On day 0 mice were anesthetized with isofluorane and intranasally infected with a dose of 10^5^ conidia (40 µL of a freshly harvested spore solution of 2.5 × 10^6^ conidia/mL). Dox was administered according to the treatment shown in Fig. S9B. Dox containing food was purchased from Envingo (Safe-diet U8200 Version 0115 A03 0.625 g/kg Doxycycline Hyclate pellets). At the selected time-point (72 h after infection for fungal burden) mice were sacrificed by a lethal injection of pentobarbital, the lungs harvested and immediately frozen.

Frozen lungs were lyophilised for 48 h in a CoolSafe ScanVac freeze drier connected to a VacuuBrand pump and subsequently ground in the presence of liquid nitrogen. DNA was isolated from the powder using the DNeasy Blood & Tissue Kit (Qiagen). DNA concentration and quality were measured using a NanoDrop 2000 (ThermoFisher Scientific). To detect the fungal burden, 500 ng of DNA extracted from each infected lung were subjected to qPCR. Primers used to amplify the *A. fumigatus* β-tubulin gene (AFUA_7G00250) were forward, 5’-ACTTCCGCAATGGACGTTAC-3’, and reverse, 5’-GGATGTTGTTGGGAATCCAC-3’. Those designed to amplify the murine actin locus (NM_007393) were forward, 5’-CGAGCACAGCTTCTTTGCAG-3’ and reverse, 5’-CCCATGGTGTCCGTTCTGA-3’. Standard curves were calculated using different concentrations of fungal and murine gDNA pure template. Negative controls containing no template DNA were subjected to the same procedure to exclude or detect any possible contamination. Three technical replicates were prepared for each lung sample. qPCRs were performed using the 7500 Fast Real-Time PCR system (Thermo Fisher Scientific) with the following thermal cycling parameters: 94 °C for 2 min and 40 cycles of 94°C for 15 s and 59°C for 1 min. The fungal burden was calculated by normalising the number of fungal genome equivalents (i.e. number of copies of the tubulin gene) to the murine genome equivalents in the sample (i.e number of copies of the actin gene) (104). Two independent experiments were carried out (*n*=9, 5 mice in the first and 4 mice in the second experiment). Burdens for each strain were compared using a Mann Whitney test.

### Molecular homology models and virtual screening

The full-length sequence for AFUA_4G07360, the cobalamin-independent methionine synthase MetH from *A. fumigatus* (AfMetH) was obtained from FungiDB (https://fungidb.org/fungidb) (105). This sequence together with the structure of the *C. albicans* orthologue (CaMetH) (PDB ID: 4L65, DOI: 10.1016/j.jmb.2014.02.006) were used to create the molecular homology model in Modeller (version 9.23) (106) with the basic option mode. The AfMetH model was then used for virtual screening with the semi-automated pipeline VSpipe (107). For comparison we also performed virtual screening with the structure of the human methionine synthase (hMS) containing the folate and homocysteine binding domains (PDB ID: 4CCZ). Docking was done using the Maybridge Ro3 1000 fragment library with AutoDock Vina (108). Results were inspected graphically using PyMol (v1.8.0.3 Enhanced for Mac OS X (Schrödinger). All images were produced with PyMol.

## ACKNOWLDGEMENTS

We acknowledge the use of the Phenotyping Center at Manchester (PCAM) for the use of their microscopes and advanced image analysis workstations. We are grateful to Prof Sven Krappmann for critical reading of the manuscript and his constant support and guidance. We also are indebted to Prof Robert Cramer for his insightful feedback on the manuscript and personal support. We would like to thank all members of MFIG for constant help and encouragement.

JA was supported by a MRC Career Development Award (MR/N008707/1). JS was supported by a BSAC scholarship (bsac-2016-0049). BPT was supported by a MRC Doctoral Training Partnership PhD studentship. Q-Exactive mass spectrometer and nanoLC instrumentation were funded by a competitive infrastructure award from Science Foundation Ireland (SFI) (12/RI/2346(3)).

## AUTHOR CONTRIBUTION

JS performed the majority of experiments, analysed and interpreted most of the data and participated in the design of the project. MS helped with the acquisition and analysis of most of the experiments. BT run the structural-based virtual screening. RAO measured SAM levels in mycelia. HMA performed the metabolomic experiment, analysed and with RG interpreted the data. RFG assisted with the mouse models of infection. DT performed the microscopy of *A. fumigatus* mitochondria and supported the other microscopy experiments. RT helped with the execution of qPCRs. KH helped to set up the GC-MS instrument. SD designed the MS/MS analysis of SAM. RG designed the metabolome analysis and interpreted the data. LT designed the virtual screening analysis and interpreted the data. EB participated in the design and conception of the project. JA conceived and designed the project and analysed most of the data.

## DATA AVAILABILITY

The raw data that support the findings of this study are available upon reasonable request to the authors. The raw data of metabolome analysis has been deposited in the MetaboLights database ^85^, under the reference MTBLS1636 (www.ebi.ac.uk/metabolights/MTBLS1636).

## SUPPLEMENTARY FIGURES

**Figure S1**.

**A)** Addition of 1 µg/mL Dox to a 16 h growing mycelia quickly downregulated transcription of *metH*. **B)** The growth of two *metH_tetOFF* strains in two different backgrounds was prevented in the presence of as little as 0.5 µg/mL Dox, even in the presence of methionine. The phenotypic analysis was repeated in three independent experiments. Representative plates are shown. **C)** An *A. fumigatus ΔmetGΔcysD* mutant (methionine auxotroph) was completely avirulent in a leukopenic murine model of invasive pulmonary aspergillosis. **D)** A strain expressing a C-terminus-GFP-tagged MetH-D616 protein from the pJA49 overexpression plasmid (Fig. S2) in the *H_OFF* background strain showed fluorescence in −Dox conditions (did not grow in +Dox conditions, Fig. S5), proving that the point mutated MetH-D616 protein is stable.

**Figure S2**.

**A)** Schematic representation of the pJA49 plasmid for episomal overexpression of genes in *A. fumigatus*. It carries the *A. nidulans* AMA1 autologous replicating sequence and the hygromycin B resistance gene (*hygrB*) as a selection marker. A unique *Stu*I restriction site allows introduction of any PCR amplified ORF in frame under the control of the *A. fumigatus* strong promoter *hspA* and the *A. nidulans trpC* terminator. **B)** Schematic representation of the genes overexpressed (in green) to eliminate potential accumulation of toxic homocysteine and derivatives (in red). **C)** Fold change in expression level of *mecA* and *sahL* measured by RT-PCR. Both genes are highly overexpressed from plasmid pJA49. **D)** Expression level of *blhA* determined by retrotranscription and PCR (fold change cannot be calculated as this gene is not expressed in *A. fumigatus*).

**Figure S3**.

Relevance of the nitrogen source for the capacity of different metabolites to reconstitute growth of the *H_OFF* strain in restrictive conditions. Amino acids alone were not able to reconstitute growth. Folic acid could partially, and ATP and SAM completely reconstituted growth when amino acids were the only N-source, suggesting that they are only taken up when there is a high variety of permeases in the cell membrane. Pyruvate could reconstitute growth to wild-type level in the absence of glucose and independently of the in presence of amino acids, although growth improved when they were added.

**Figure S4**.

**A)** Principal component analysis (PCA) scores plot of the GC-MS metabolome analysis. The wild-type and *H_OFF* strains clusters were close before addition of Dox and became clearly separated upon 6 h incubation in the presence of Dox. **B)** The content of several amino acids and sugars were reduced in the H_OFF+Dox sample compared with the developmentally matched wt+Dox sample. **C)** Neither low concentrations of rapamycin (to partially inhibit TOR) nor of H89 (to partially inhibit PKA) were able to reconstitute growth of the *H_OFF* strain in restrictive conditions. This suggests that these regulatory pathways are not responsible for the deleterious switch in metabolism that causes a drop in cellular ATP.

**Figure S5**.

**A)** Expression of MetH-GFP wild type and MetH^R742A^-GFP reconstituted growth of the *H_OFF* strain in restrictive conditions, demonstrating that the proteins are active. As expected, the inactive protein MetH^D616A^-GFP did not trigger growth. **B)** Western-blot of MetH. Strains expressing MetH-GFP wild type and MetH^R742A^-GFP were grown in the presence of Dox, nuclei isolated from the mycelia (as described in material and methods), proteins purified and blotted with an anti-GFP antibody. The MetH^R742A^-GFP protein could be detected in nuclei, at similar levels as the wild-type MetH-GFP.

**Figure S6**.

Doxycycline can perturb *A. fumigatus* hyphal mitochondrial morphology at high concentrations. *A. fumigatus* hyphae were stained with 10 µM Rhodamine 123 after overnight incubation in **A)** 0 µg/ml, **B)** 1000 µg/ml, **C)** 1 µg/ml, **D)** 10 µg/ml and **E)** 100 µg/ml Doxycycline. Representative single plane confocal fluorescence images are displayed in A-E. Notice the granulation in B (1000 µg/ml and variable response in E (100 µg/ml), whereas C (1 µg/ml) and D (10 µg/ml) display normal mitochondrial morphology. Scale bars = 10 µm.

**Figure S7**.

**A)** Measurement of fungal growth by Optical Density (OD) showed that addition of 1 µg/mL Dox 12 or 16 h after inoculation significantly reduced fugal growth at 24 and 36 h post-incubation. **B)** Microscopy analysis of the effect of the addition of 2 concentrations of Dox to resting or 8 h germinated conidia of the *H_OFF* strain. In the absence of methionine growth is immediately halted in a sustained manner. In the presence of methionine conidia can germinate and germlings elongate the growing tube before growth stops. However, once arrested, growth is halted in a sustained manner. Microimages were taken with a wide-field microscope at 16 h post-inoculation and with a stereomicroscope 40 h after inoculation (to be able to capture the huge mass of fungal growth in the control conditions).

**Figure S8**.

**A)** Colony formation of the strain *cyp51A_tetOFFΔcyp51B* (*51A_OFF*) is completely prevented with as little as 0.1 µg/mL on an agar plate. Phenotypic analysis was repeated in two independent experiments. **B)** Microscopy analysis of the effect of the addition of 2 concentrations of Dox to resting or 8 h germinated conidia of the 51A*_OFF* strain. In both cases growth is immediately blocked. We could observe bursting germlings with high doses of Dox (red arrow).

**Figure S9.**

**A)** Dox regime applied to *Galleria mellonella*. A maximum of 5 doses (50 mg/kg) were applied, commencing either at the time of infection or 6 h later. **B)** Dox regimen administered to mice. Treatment started 16 h after infection with a subcutaneous (SC) Dox injection (50 mg/kg) and change to Dox food. Treatment was maintained with SC injections every 12 h after the infection time-point. Dox concentration in the lungs was measured in a preliminary experiment with uninfected mice. Lungs were harvested 4 h after the beginning of treatment, 2 h after the third injection (on day 2) and 9 h after the 5^th^ (on day 3, last injection). The concentrations were determined to range from 0.9 to 2.2 µg/mL, sufficient to downregulate gene expression according to the results *in vitro*.

**Video S1**

Time-lapse of growth of the *H_OFF* strain upon Dox addition and withdrawal.

**Video S2**

Time-lapse of growth of the *wild-type* strains upon Dox addition.

## REFERENCES

1. Fisher MC, Hawkins NJ, Sanglard D, Gurr SJ. 2018. Worldwide emergence of resistance to antifungal drugs challenges human health and food security. Science 360:739–742.

2. Bongomin F, Gago S, Oladele RO, Denning DW. 2017. Global and Multi-National Prevalence of Fungal Diseases-Estimate Precision. J Fungi (Basel) 3.

3. Bitar D, Lortholary O, Le Strat Y, Nicolau J, Coignard B, Tattevin P, Che D, Dromer F. 2014. Population-based analysis of invasive fungal infections, France, 2001-2010. Emerg Infect Dis 20:1149–55.

4. Brown GD, Denning DW, Gow NA, Levitz SM, Netea MG, White TC. 2012. Hidden killers: human fungal infections. Sci Transl Med 4:165rv13.

5. Nyazika TK, Hagen F, Machiridza T, Kutepa M, Masanganise F, Hendrickx M, Boekhout T, Magombei-Majinjiwa T, Siziba N, Chin’ombe N, Mateveke K, Meis JF, Robertson VJ. 2016. *Cryptococcus neoformans* population diversity and clinical outcomes of HIV-associated cryptococcal meningitis patients in Zimbabwe. J Med Microbiol 65:1281–1288.

6. Patterson TF, Thompson GR, 3rd, Denning DW, Fishman JA, Hadley S, Herbrecht R, Kontoyiannis DP, Marr KA, Morrison VA, Nguyen MH, Segal BH, Steinbach WJ, Stevens DA, Walsh TJ, Wingard JR, Young JA, Bennett JE. 2016. Executive Summary: Practice Guidelines for the Diagnosis and Management of Aspergillosis: 2016 Update by the Infectious Diseases Society of America. Clin Infect Dis 63:433–42.

7. Pound MW, Townsend ML, Dimondi V, Wilson D, Drew RH. 2011. Overview of treatment options for invasive fungal infections. Med Mycol 49:561–80.

8. Ghannoum MA, Rice LB. 1999. Antifungal agents: mode of action, mechanisms of resistance, and correlation of these mechanisms with bacterial resistance. Clin Microbiol Rev 12:501–17.

9. Verweij PE, Chowdhary A, Melchers WJ, Meis JF. 2016. Azole Resistance in *Aspergillus fumigatus*: Can We Retain the Clinical Use of Mold-Active Antifungal Azoles? Clin Infect Dis 62:362–8.

10. Amich J, Krappmann S. 2012. Deciphering metabolic traits of the fungal pathogen *Aspergillus fumigatus*: redundancy vs. essentiality. Frontiers in Microbiology Fungi and Their Interactions.

11. Kaltdorf M, Srivastava M, Gupta SK, Liang C, Binder J, Dietl AM, Meir Z, Haas H, Osherov N, Krappmann S, Dandekar T. 2016. Systematic Identification of Anti-Fungal Drug Targets by a Metabolic Network Approach. Front Mol Biosci 3:22.

12. Oliver JD, Sibley GEM, Beckmann N, Dobb KS, Slater MJ, McEntee L, du Pre S, Livermore J, Bromley MJ, Wiederhold NP, Hope WW, Kennedy AJ, Law D, Birch M. 2016. F901318 represents a novel class of antifungal drug that inhibits dihydroorotate dehydrogenase. Proc Natl Acad Sci U S A 113:12809–12814.

13. Evans JC, Huddler DP, Hilgers MT, Romanchuk G, Matthews RG, Ludwig ML. 2004. Structures of the N-terminal modules imply large domain motions during catalysis by methionine synthase. Proceedings of the National Academy of Sciences of the United States of America 101:3729–3736.

14. Peariso K, Zhou ZHS, Smith AE, Matthews RG, Penner-Hahn JE. 2001. Characterization of the zinc sites in cobalamin-independent and cobalamin-dependent methionine synthase using zinc and selenium X-ray absorption spectroscopy. Biochemistry 40:987–993.

15. Gonzalez JC, Banerjee RV, Huang S, Sumner JS, Matthews RG. 1992. Comparison of cobalamin-independent and cobalamin-dependent methionine synthases from *Escherichia coli:* two solutions to the same chemical problem. Biochemistry 31:6045–56.

16. Matthews RG, Smith AE, Zhou ZHS, Taurog RE, Bandarian V, Evans JC, Ludwig M. 2003. Cobalamin-dependent and cobalamin-independent methionine synthases: Are there two solutions to the same chemical problem? Helvetica Chimica Acta 86:3939–3954.

17. Amich J, Dumig M, O’Keeffe G, Binder J, Doyle S, Beilhack A, Krappmann S. 2016. Exploration of Sulfur Assimilation of *Aspergillus fumigatus* Reveals Biosynthesis of Sulfur-Containing Amino Acids as a Virulence Determinant. Infect Immun 84:917–29.

18. Segal ES, Gritsenko V, Levitan A, Yadav B, Dror N, Steenwyk JL, Silberberg Y, Mielich K, Rokas A, Gow NAR, Kunze R, Sharan R, Berman J. 2018. Gene Essentiality Analyzed by In Vivo Transposon Mutagenesis and Machine Learning in a Stable Haploid Isolate of *Candida albicans*. MBio 9.

19. Suliman HS, Appling DR, Robertus JD. 2007. The gene for cobalamin-independent methionine synthase is essential in *Candida albicans*: a potential antifungal target. Arch Biochem Biophys 467:218–26.

20. Pascon RC, Ganous TM, Kingsbury JM, Cox GM, McCusker JH. 2004. *Cryptococcus neoformans* methionine synthase: expression analysis and requirement for virulence. Microbiology 150:3013–23.

21. Li W, Cowley A, Uludag M, Gur T, McWilliam H, Squizzato S, Park YM, Buso N, Lopez R. 2015. The EMBL-EBI bioinformatics web and programmatic tools framework. Nucleic Acids Res 43:W580–4.

22. McWilliam H, Li W, Uludag M, Squizzato S, Park YM, Buso N, Cowley AP, Lopez R. 2013. Analysis Tool Web Services from the EMBL-EBI. Nucleic Acids Res 41:W597–600.

23. Miwa GT. 2010. The Drug Discovery Process p1–14. In Lu C, Li AP (ed), Enzyme Inhibition in Drug Discovery and Development: The Good and the Bad doi:10.1002/9780470538951.

24. Cook D, Brown D, Alexander R, March R, Morgan P, Satterthwaite G, Pangalos MN. 2014. Lessons learned from the fate of AstraZeneca’s drug pipeline: a five-dimensional framework. Nature Reviews Drug Discovery 13:419–431.

25. Wanka F, Cairns T, Boecker S, Berens C, Happel A, Zheng X, Sun J, Krappmann S, Meyer V. 2016. Tet-on, or Tet-off, that is the question: Advanced conditional gene expression in *Aspergillus*. Fungal Genet Biol 89:72–83.

26. Scott J, Sueiro-Olivares M, Ahmed W, Heddergott C, Zhao C, Thomas R, Bromley M, Latge JP, Krappmann S, Fowler S, Bignell E, Amich J. 2019. *Pseudomonas aeruginosa*-Derived Volatile Sulfur Compounds Promote Distal *Aspergillus fumigatus* Growth and a Synergistic Pathogen-Pathogen Interaction That Increases Pathogenicity in Co-infection. Front Microbiol 10:2311.

27. Lucock M. 2000. Folic acid: nutritional biochemistry, molecular biology, and role in disease processes. Mol Genet Metab 71:121–38.

28. Tzin V, Galili G. 2010. The Biosynthetic Pathways for Shikimate and Aromatic Amino Acids in *Arabidopsis thaliana*. Arabidopsis Book 8:e0132.

29. Deakova Z, Durackova Z, Armstrong DW, Lehotay J. 2015. Two-dimensional high performance liquid chromatography for determination of homocysteine, methionine and cysteine enantiomers in human serum. J Chromatogr A 1408:118–24.

30. Elshorbagy AK, Jerneren F, Samocha-Bonet D, Refsum H, Heilbronn LK. 2016. Serum S-adenosylmethionine, but not methionine, increases in response to overfeeding in humans. Nutr Diabetes 6:e192.

31. Jochim A, Shi T, Belikova D, Schwarz S, Peschel A, Heilbronner S. 2019. Methionine Limitation Impairs Pathogen Expansion and Biofilm Formation Capacity. Appl Environ Microbiol 85.

32. Ubhi D, Kago G, Monzingo AF, Robertus JD. 2014. Structural analysis of a fungal methionine synthase with substrates and inhibitors. J Mol Biol 426:1839–47.

33. Sahu U, Rajendra VKH, Kapnoor SS, Bhagavat R, Chandra N, Rangarajan PN. 2017. Methionine synthase is localized to the nucleus in *Pichia pastoris* and *Candida albicans* and to the cytoplasm in *Saccharomyces cerevisiae*. J Biol Chem 292:14730–14746.

34. Fujita Y, Ukena E, Iefuji H, Giga-Hama Y, Takegawa K. 2006. Homocysteine accumulation causes a defect in purine biosynthesis: further characterization of *Schizosaccharomyces pombe* methionine auxotrophs. Microbiology 152:397–404.

35. Gems D, Johnstone IL, Clutterbuck AJ. 1991. An autonomously replicating plasmid transforms *Aspergillus nidulans* at high frequency. Gene 98:61–7.

36. Aleksenko AY, Clutterbuck AJ. 1995. Recombinational stability of replicating plasmids in *Aspergillus nidulans* during transformation, vegetative growth and sexual reproduction. Curr Genet 28:87–93.

37. Paul S, Klutts JS, Moye-Rowley WS. 2012. Analysis of promoter function in *Aspergillus fumigatus*. Eukaryot Cell 11:1167–77.

38. Christopher SA, Melnyk S, James SJ, Kruger WD. 2002. S-adenosylhomocysteine, but not homocysteine, is toxic to yeast lacking cystathionine beta-synthase. Mol Genet Metab 75:335–43.

39. Perna AF, Ingrosso D, Lombardi C, Acanfora F, Satta E, Cesare CM, Violetti E, Romano MM, De Santo NG. 2003. Possible mechanisms of homocysteine toxicity. Kidney Int Suppl doi:10.1046/j.1523-1755.63.s84.33.x:S137-40.

40. Bretes E, Zimny J. 2013. Homocysteine thiolactone affects protein ubiquitination in yeast. Acta Biochim Pol 60:485–8.

41. Jakubowski H. 2006. Pathophysiological consequences of homocysteine excess. J Nutr 136:1741S–1749S.

42. Lopez-Ibanez J, Pazos F, Chagoyen M. 2016. MBROLE 2.0-functional enrichment of chemical compounds. Nucleic Acids Research 44:W201–W204.

43. Chong J, Soufan O, Li C, Caraus I, Li SZ, Bourque G, Wishart DS, Xia JG. 2018. MetaboAnalyst 4.0: towards more transparent and integrative metabolomics analysis. Nucleic Acids Research 46:W486–W494.

44. Xia JG, Psychogios N, Young N, Wishart DS. 2009. MetaboAnalyst: a web server for metabolomic data analysis and interpretation. Nucleic Acids Research 37:W652–W660.

45. Fendt SM, Sauer U. 2010. Transcriptional regulation of respiration in yeast metabolizing differently repressive carbon substrates. BMC Syst Biol 4:12.

46. Akita O, Nishimori C, Shimamoto T, Fujii T, Iefuji H. 2000. Transport of pyruvate in *Saccharomyces cerevisiae* and cloning of the gene encoded pyruvate permease. Biosci Biotechnol Biochem 64:980–4.

47. Forte GM, Davie E, Lie S, Franz-Wachtel M, Ovens AJ, Wang T, Oakhill JS, Macek B, Hagan IM, Petersen J. 2019. Import of extracellular ATP in yeast and man modulates AMPK and TORC1 signalling. J Cell Sci 132.

48. Hu W, Sillaots S, Lemieux S, Davison J, Kauffman S, Breton A, Linteau A, Xin C, Bowman J, Becker J, Jiang B, Roemer T. 2007. Essential gene identification and drug target prioritization in *Aspergillus fumigatus*. PLoS Pathog 3:e24.

49. Beattie SR, Mark KMK, Thammahong A, Ries LNA, Dhingra S, Caffrey-Carr AK, Cheng C, Black CC, Bowyer P, Bromley MJ, Obar JJ, Goldman GH, Cramer RA. 2017. Filamentous fungal carbon catabolite repression supports metabolic plasticity and stress responses essential for disease progression. PLoS Pathog 13:e1006340.

50. Cramer RA. 2016. *In vivo* veritas: *Aspergillus fumigatus* proliferation and pathogenesis--conditionally speaking. Virulence 7:7–10.

51. Kowalski CH, Kerkaert JD, Liu KW, Bond MC, Hartmann R, Nadell CD, Stajich JE, Cramer RA. 2019. Fungal biofilm morphology impacts hypoxia fitness and disease progression. Nat Microbiol 4:2430–2441.

52. Gorman MW, Feigl EO, Buffington CW. 2007. Human plasma ATP concentration. Clin Chem 53:318–25.

53. Ito S, Furuya K, Sokabe M, Hasegawa Y. 2016. Cellular ATP release in the lung and airway. Aims Biophysics 3:571–584.

54. Mortaz E, Braber S, Nazary M, Givi ME, Nijkamp FP, Folkerts G. 2009. ATP in the pathogenesis of lung emphysema. Eur J Pharmacol 619:92–6.

55. Riteau N, Gasse P, Fauconnier L, Gombault A, Couegnat M, Fick L, Kanellopoulos J, Quesniaux VFJ, Marchand-Adam S, Crestani B, Ryffel B, Couillin I. 2010. Extracellular ATP Is a Danger Signal Activating P2X(7) Receptor in Lung Inflammation and Fibrosis. American Journal of Respiratory and Critical Care Medicine 182:774–783.

56. Walvekar AS, Srinivasan R, Gupta R, Laxman S. 2018. Methionine coordinates a hierarchically organized anabolic program enabling proliferation. Mol Biol Cell 29:3183–3200.

57. De Virgilio C, Loewith R. 2006. The TOR signalling network from yeast to man. Int J Biochem Cell Biol 38:1476–81.

58. Loewith R, Hall MN. 2011. Target of rapamycin (TOR) in nutrient signaling and growth control. Genetics 189:1177–201.

59. Saxton RA, Sabatini DM. 2017. mTOR Signaling in Growth, Metabolism, and Disease. Cell 168:960–976.

60. Thiepold AL, Lorenz NI, Foltyn M, Engel AL, Dive I, Urban H, Heller S, Bruns I, Hofmann U, Drose S, Harter PN, Mittelbronn M, Steinbach JP, Ronellenfitsch MW. 2017. Mammalian target of rapamycin complex 1 activation sensitizes human glioma cells to hypoxia-induced cell death. Brain 140:2623–2638.

61. Tsouko E, Khan AS, White MA, Han JJ, Shi Y, Merchant FA, Sharpe MA, Xin L, Frigo DE. 2014. Regulation of the pentose phosphate pathway by an androgen receptor-mTOR-mediated mechanism and its role in prostate cancer cell growth. Oncogenesis 3:e103.

62. Baldin C, Valiante V, Kruger T, Schafferer L, Haas H, Kniemeyer O, Brakhage AA. 2015. Comparative proteomics of a tor inducible *Aspergillus fumigatus* mutant reveals involvement of the Tor kinase in iron regulation. Proteomics 15:2230–43.

63. Caza M, Kronstad JW. 2019. The cAMP/Protein Kinase a Pathway Regulates Virulence and Adaptation to Host Conditions in *Cryptococcus neoformans*. Front Cell Infect Microbiol 9:212.

64. Grosse C, Heinekamp T, Kniemeyer O, Gehrke A, Brakhage AA. 2008. Protein kinase A regulates growth, sporulation, and pigment formation in *Aspergillus fumigatus*. Appl Environ Microbiol 74:4923–33.

65. Gerke J, Bayram O, Braus GH. 2012. Fungal S-adenosylmethionine synthetase and the control of development and secondary metabolism in *Aspergillus nidulans*. Fungal Genet Biol 49:443–54.

66. Moullan N, Mouchiroud L, Wang X, Ryu D, Williams EG, Mottis A, Jovaisaite V, Frochaux MV, Quiros PM, Deplancke B, Houtkooper RH, Auwerx J. 2015. Tetracyclines Disturb Mitochondrial Function across Eukaryotic Models: A Call for Caution in Biomedical Research. Cell Rep 10:1681–1691.

67. Samant R, Ahler E, Sullivan WJ, Cass A, Braas D, York AG, Bensinger SJ, Graeber TG, Christofk HR. 2013. Doxycycline Alters Metabolism and Proliferation of Human Cell Lines. PLoS ONE 8.

68. Sasse A, Hamer SN, Amich J, Binder J, Krappmann S. 2015. Mutant characterization and in vivo conditional repression identify aromatic amino acid biosynthesis to be essential for *Aspergillus fumigatus* virulence. Virulence 7:56–62.

69. Binder J, Shadkchan Y, Osherov N, Krappmann S. 2020. The Essential Thioredoxin Reductase of the Human Pathogenic Mold *Aspergillus fumigatus* Is a Promising Antifungal Target. Frontiers in Microbiology 11.

70. Miceli MH, Bernardo SM, Lee SA. 2009. In vitro analyses of the combination of high-dose doxycycline and antifungal agents against *Candida albicans* biofilms. International Journal of Antimicrobial Agents 34:326–332.

71. Chen T, Wagner AS, Tams RN, Eyer JE, Kauffman SJ, Gann ER, Fernandez EJ, Reynolds TB, Heitman J. 2019. Lrg1 Regulates β (1,3)-Glucan Masking in *Candida albicans* through the Cek1 MAP Kinase Pathway. mBio 10.

72. Stewart JIP, Fava VM, Kerkaert JD, Subramanian AS, Gravelat FN, Lehoux M, Howell PL, Cramer RA, Sheppard DC. 2020. Reducing *Aspergillus fumigatus* Virulence through Targeted Dysregulation of the Conidiation Pathway. mBio 11.

73. Fiori A, Van Dijck P. 2012. Potent Synergistic Effect of Doxycycline with Fluconazole against *Candida albicans* Is Mediated by Interference with Iron Homeostasis. Antimicrobial Agents and Chemotherapy 56:3785–3796.

74. Oliver BG, Silver PM, Marie C, Hoot SJ, Leyde SE, White TC. 2008. Tetracycline alters drug susceptibility in *Candida albicans* and other pathogenic fungi. Microbiology 154:960–970.

75. Wishart JA, Hayes A, Wardleworth L, Zhang N, Oliver SG. 2005. Doxycycline, the drug used to control thetet-regulatable promoter system, has no effect on global gene expression in *Saccharomyces cerevisiae*. Yeast 22:565–569.

76. Xie JL, Bohovych I, Wong EOY, Lambert J-P, Gingras A-C, Khalimonchuk O, Cowen LE, Leach MD. 2017. Ydj1 governs fungal morphogenesis and stress response, and facilitates mitochondrial protein import via Mas1 and Mas2. Microbial Cell 4:342–361.

77. Bauer I, Misslinger M, Shadkchan Y, Dietl AM, Petzer V, Orasch T, Abt B, Graessle S, Osherov N, Haas H. 2019. The Lysine Deacetylase RpdA Is Essential for Virulence in *Aspergillus fumigatus*. Front Microbiol 10:2773.

78. Grenier D, Huot MP, Mayrand D. 2000. Iron-chelating activity of tetracyclines and its impact on the susceptibility of *Actinobacillus actinomycetemcomitans* to these antibiotics. Antimicrob Agents Chemother 44:763–6.

79. Peng Y, Zhang H, Xu M, Tan MW. 2018. A Tet-Off gene expression system for validation of antifungal drug targets in a murine invasive pulmonary aspergillosis model. Sci Rep 8:443.

80. Ferrer JL, Ravanel S, Robert M, Dumas R. 2004. Crystal structures of cobalamin-independent methionine synthase complexed with zinc, homocysteine, and methyltetrahydrofolate. J Biol Chem 279:44235–8.

81. Warnock DW, Johnson EM, Burke J, Pracharktam R. 1989. Effect of methotrexate alone and in combination with antifungal drugs on the growth of *Candida albicans*. J Antimicrob Chemother 23:837–47.

82. Yang J, Gao L, Yu P, Kosgey JC, Jia L, Fang Y, Xiong J, Zhang F. 2019. In vitro synergy of azole antifungals and methotrexate against *Candida albicans*. Life Sci 235:116827.

83. Fan CC, Vitols KS, Huennekens FM. 1980. Inhibition of dihydrofolate reductase by methotrexate: a new look at an old problem. Adv Enzyme Regul 18:41–52.

84. Woodcock DM, Crowther PJ, Doherty J, Jefferson S, DeCruz E, Noyer-Weidner M, Smith SS, Michael MZ, Graham MW. 1989. Quantitative evaluation of *Escherichia coli* host strains for tolerance to cytosine methylation in plasmid and phage recombinants. Nucleic Acids Res 17:3469–78.

85. Amich J, Schafferer L, Haas H, Krappmann S. 2013. Regulation of sulphur assimilation is essential for virulence and affects iron homeostasis of the human-pathogenic mould *Aspergillus fumigatus*. PLoS Pathog 9:e1003573.

86. Wood WB. 1966. Host specificity of DNA produced by *Escherichia coli*: bacterial mutations affecting the restriction and modification of DNA. J Mol Biol 16:118–33.

87. da Silva Ferreira ME, Kress MR, Savoldi M, Goldman MH, Hartl A, Heinekamp T, Brakhage AA, Goldman GH. 2006. The akuB(KU80) mutant deficient for nonhomologous end joining is a powerful tool for analyzing pathogenicity in *Aspergillus fumigatus*. Eukaryot Cell 5:207–11.

88. Szewczyk E, Nayak T, Oakley CE, Edgerton H, Xiong Y, Taheri-Talesh N, Osmani SA, Oakley BR. 2006. Fusion PCR and gene targeting in *Aspergillus nidulans*. Nat Protoc 1:3111–20.

89. Kafer E. 1977. Meiotic and mitotic recombination in *Aspergillus* and its chromosomal aberrations. Adv Genet 19:33–131.

90. Sambrook J, Fritsch EF, Maniatis T. 1989. Molecular Cloning: A Laboratory Manual, NY.

91. Kolar M, Punt PJ, van den Hondel CA, Schwab H. 1988. Transformation of *Penicillium chrysogenum* using dominant selection markers and expression of an *Escherichia coli lacZ* fusion gene. Gene 62:127–34.

92. Southern E. 2006. Southern blotting. Nat Protoc 1:518–25.

93. Southern EM. 1975. Detection of specific sequences among DNA fragments separated by gel electrophoresis. J Mol Biol 98:503–17.

94. Reverberi M, Punelli M, Smith CA, Zjalic S, Scarpari M, Scala V, Cardinali G, Aspite N, Pinzari F, Payne GA, Fabbri AA, Fanelli C. 2012. How peroxisomes affect aflatoxin biosynthesis in *Aspergillus flavus*. PLoS One 7:e48097.

95. Wedge DC, Allwood JW, Dunn W, Vaughan AA, Simpson K, Brown M, Priest L, Blackhall FH, Whetton AD, Dive C, Goodacre R. 2011. Is Serum or Plasma More Appropriate for Intersubject Comparisons in Metabolomic Studies? An Assessment in Patients with Small-Cell Lung Cancer. Analytical Chemistry 83:6689–6697.

96. Sumner LW, Amberg A, Barrett D, Beale MH, Beger R, Daykin CA, Fan TWM, Fiehn O, Goodacre R, Griffin JL, Hankemeier T, Hardy N, Harnly J, Higashi R, Kopka J, Lane AN, Lindon JC, Marriott P, Nicholls AW, Reily MD, Thaden JJ, Viant MR. 2007. Proposed minimum reporting standards for chemical analysis. Metabolomics 3:211–221.

97. Gromski PS, Muhamadali H, Ellis DI, Xu Y, Correa E, Turner ML, Goodacre R. 2015. A tutorial review: Metabolomics and partial least squares-discriminant analysis - a marriage of convenience or a shotgun wedding. Analytica Chimica Acta 879:10–23.

98. Haug K, Cochrane K, Nainala VC, Williams M, Chang J, Jayaseelan KV, O’Donovan C. 2020. MetaboLights: a resource evolving in response to the needs of its scientific community. Nucleic Acids Res 48:D440–D444.

99. Owens RA, O’Keeffe G, Smith EB, Dolan SK, Hammel S, Sheridan KJ, Fitzpatrick DA, Keane TM, Jones GW, Doyle S. 2015. Interplay between Gliotoxin Resistance, Secretion, and the Methyl/Methionine Cycle in *Aspergillus fumigatus*. Eukaryot Cell 14:941–57.

100. Sperling AS, Grunstein M. 2009. Histone H3 N-terminus regulates higher order structure of yeast heterochromatin. Proc Natl Acad Sci U S A 106:13153–9.

101. Schindelin J, Arganda-Carreras I, Frise E, Kaynig V, Longair M, Pietzsch T, Preibisch S, Rueden C, Saalfeld S, Schmid B, Tinevez JY, White DJ, Hartenstein V, Eliceiri K, Tomancak P, Cardona A. 2012. Fiji: an open-source platform for biological-image analysis. Nature Methods 9:676–682.

102. Mutterer J, Zinck E. 2013. Quick-and-clean article figures with FigureJ. Journal of Microscopy 252:89–91.

103. Kavanagh K, Fallon JP. 2010. *Galleria mellonella* larvae as models for studying fungal virulence. Fungal Biol Rev 24:79–83.

104. Gago S, Buitrago MJ, Clemons KV, Cuenca-Estrella M, Mirels LF, Stevens DA. 2014. Development and validation of a quantitative real-time PCR assay for the early diagnosis of coccidioidomycosis. Diagn Microbiol Infect Dis 79:214–21.

105. Stajich JE, Harris T, Brunk BP, Brestelli J, Fischer S, Harb OS, Kissinger JC, Li W, Nayak V, Pinney DF, Stoeckert CJ, Jr., Roos DS. 2012. FungiDB: an integrated functional genomics database for fungi. Nucleic Acids Res 40:D675–81.

106. Fiser A, Sali A. 2003. MODELLER: Generation and refinement of homology-based protein structure models. Macromolecular Crystallography, Pt D 374:461–491.

107. Alvarez-Carretero S, Pavlopoulou N, Adams J, Gilsenan J, Tabernero L. 2018. VSpipe, an Integrated Resource for Virtual Screening and Hit Selection: Applications to Protein Tyrosine Phospahatase Inhibition. Molecules 23.

108. Trott O, Olson AJ. 2010. Software News and Update AutoDock Vina: Improving the Speed and Accuracy of Docking with a New Scoring Function, Efficient Optimization, and Multithreading. Journal of Computational Chemistry 31:455–461.

