## Supplementary figures and images for "Targeting methionine synthase in a fungal pathogen causes a metabolic imbalance that impacts cell energetics, growth and virulence"

### Supplementary figures

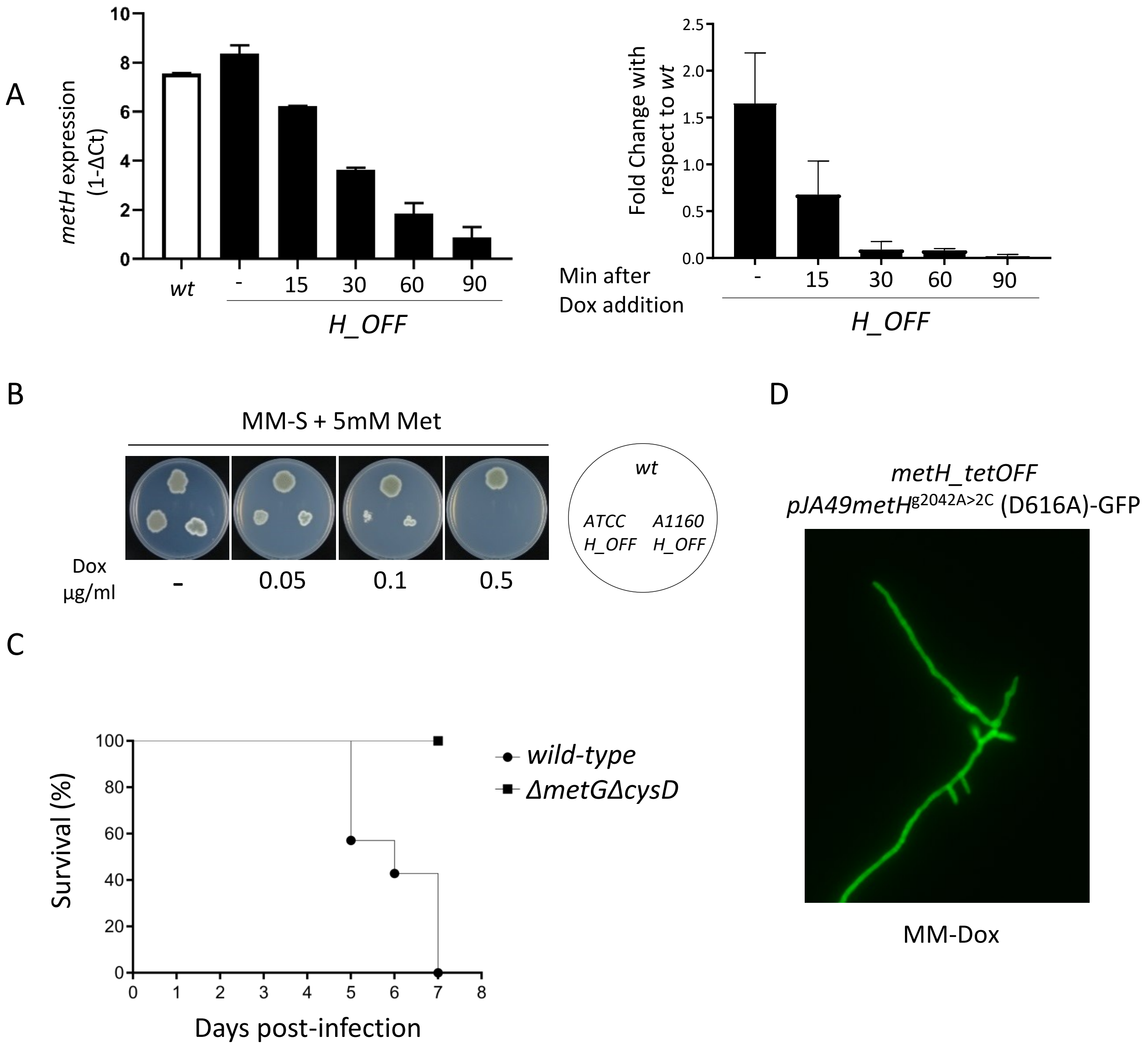

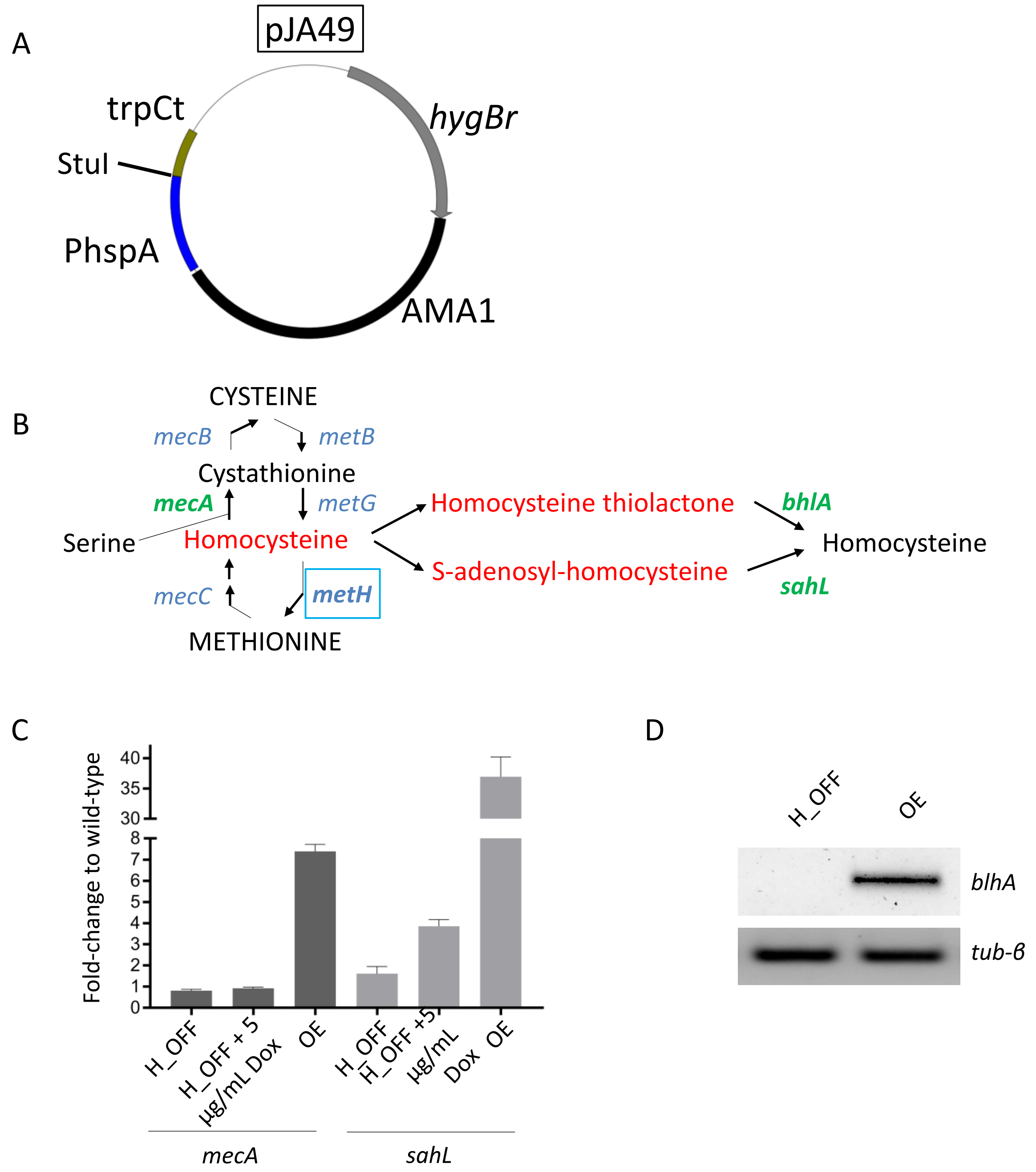

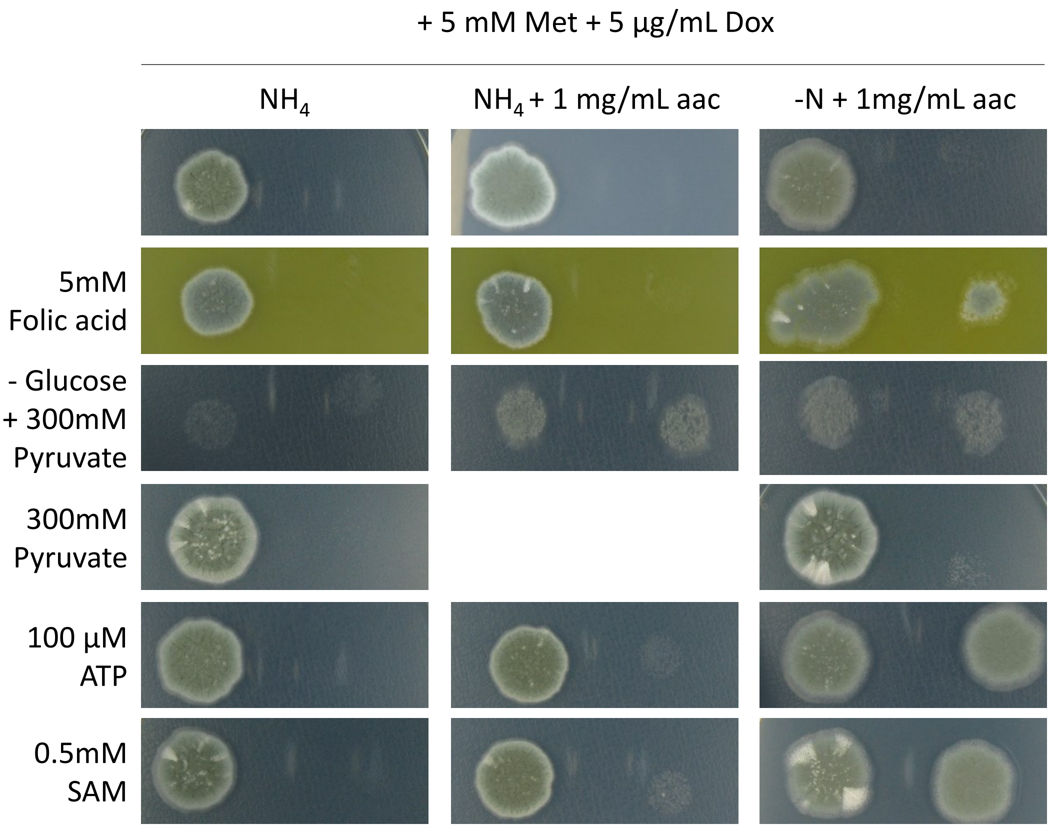

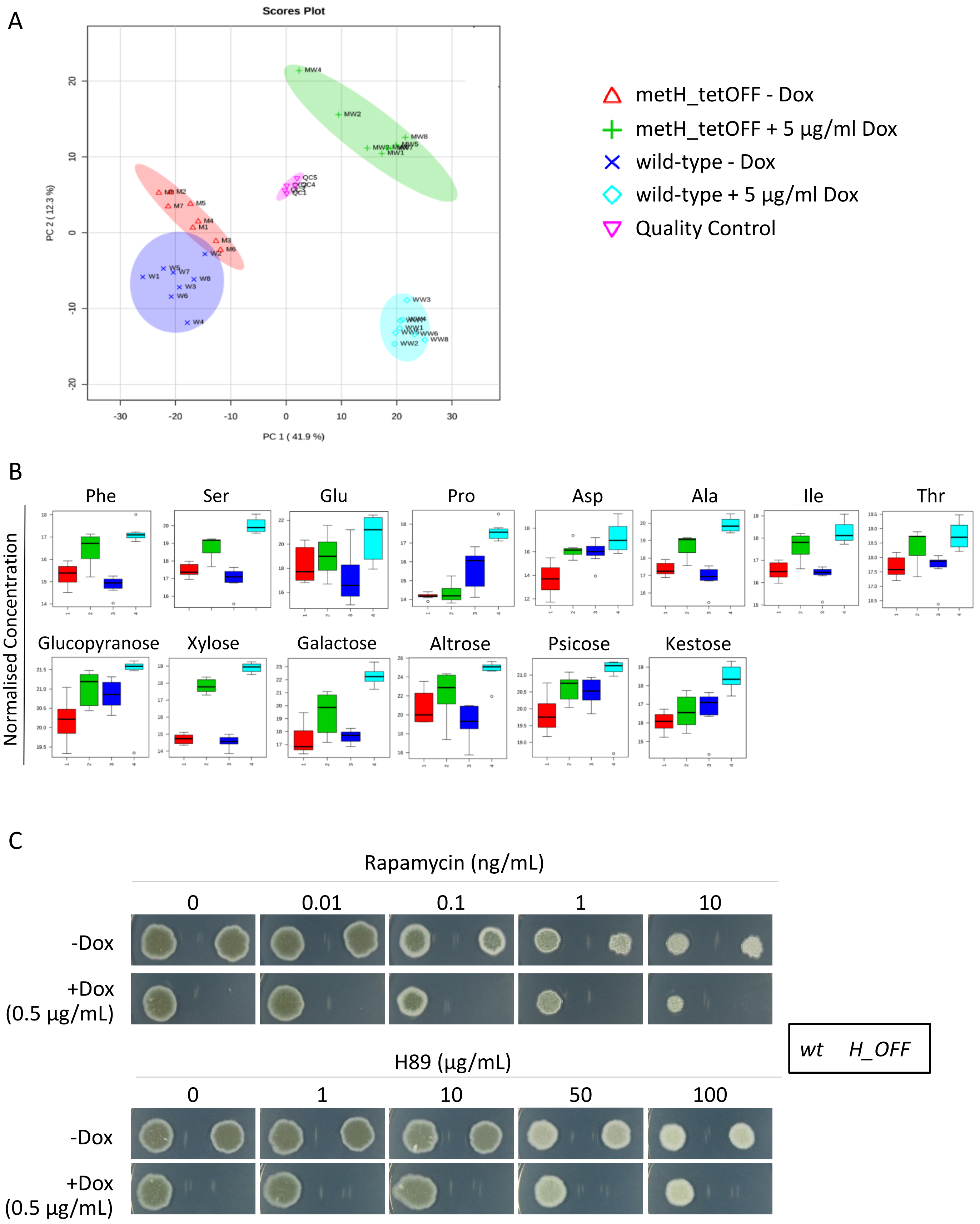

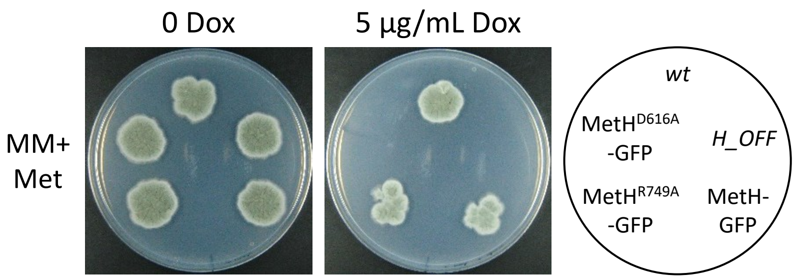

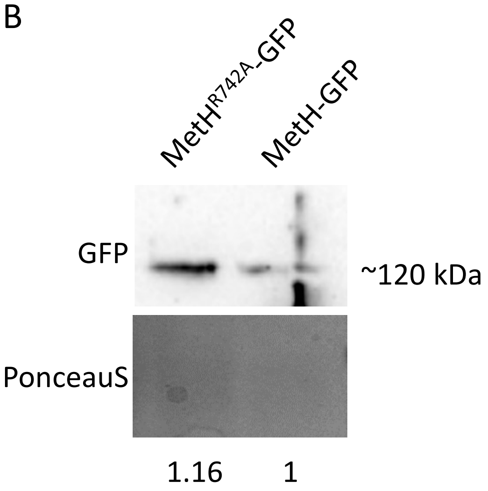

Fig. S6

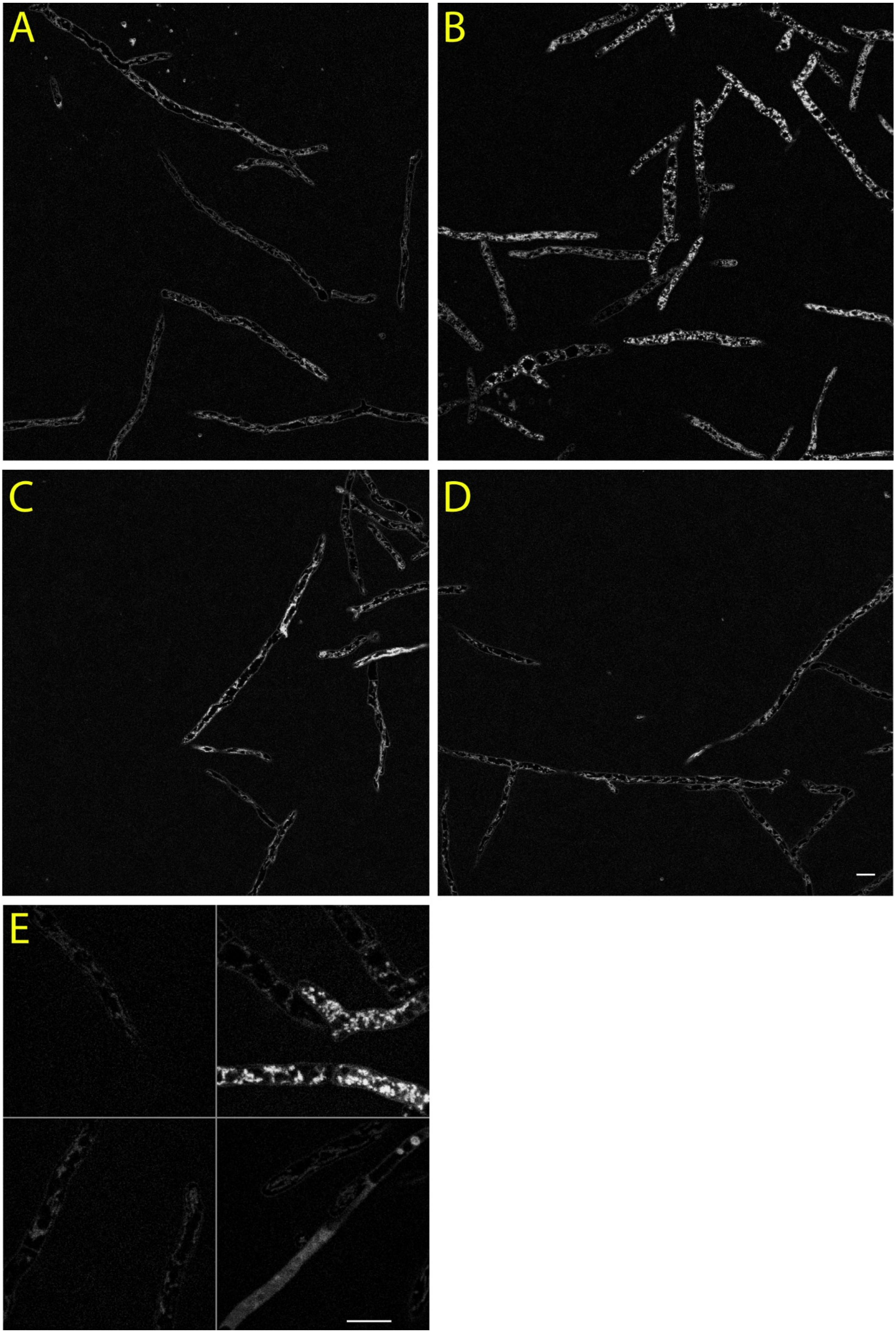

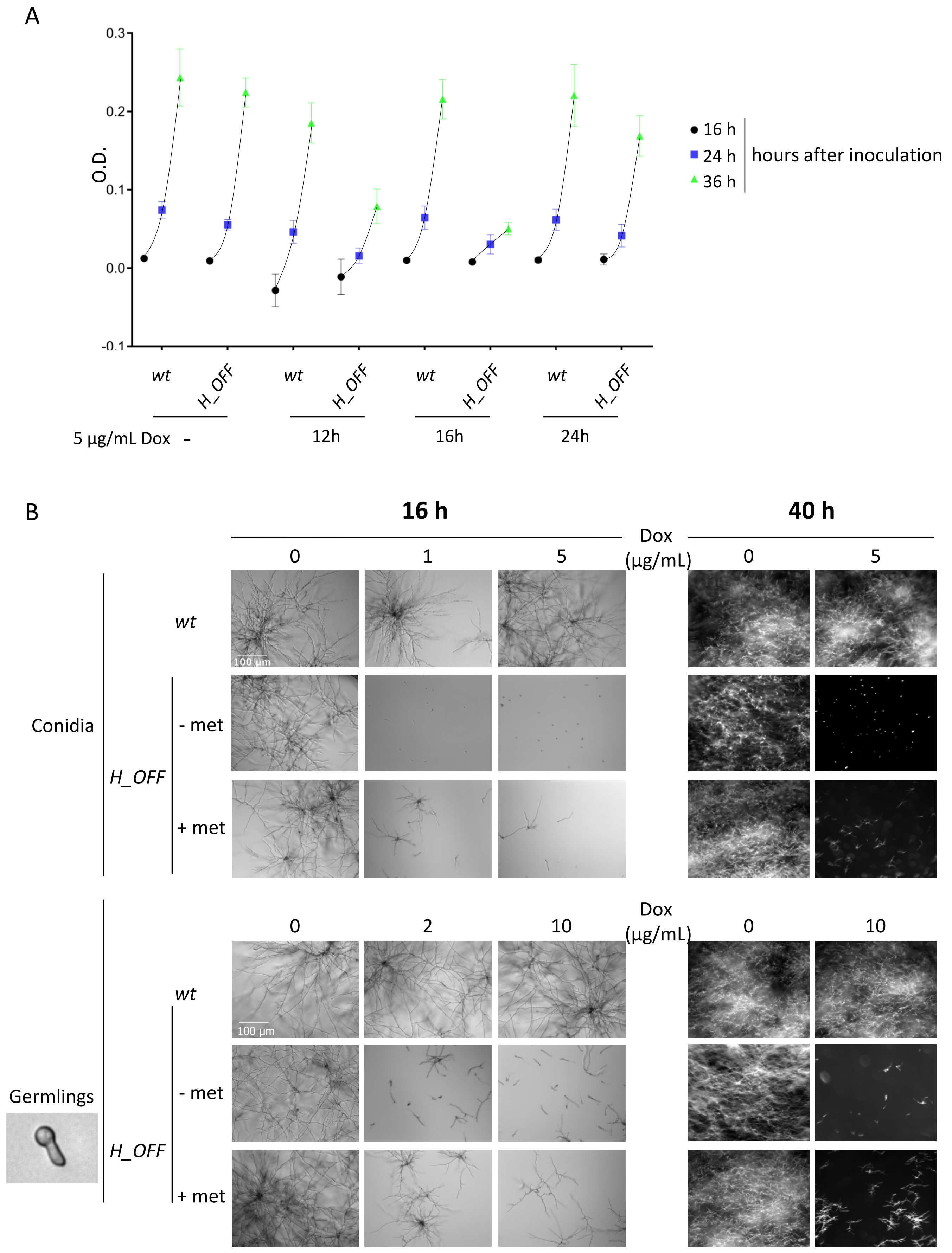

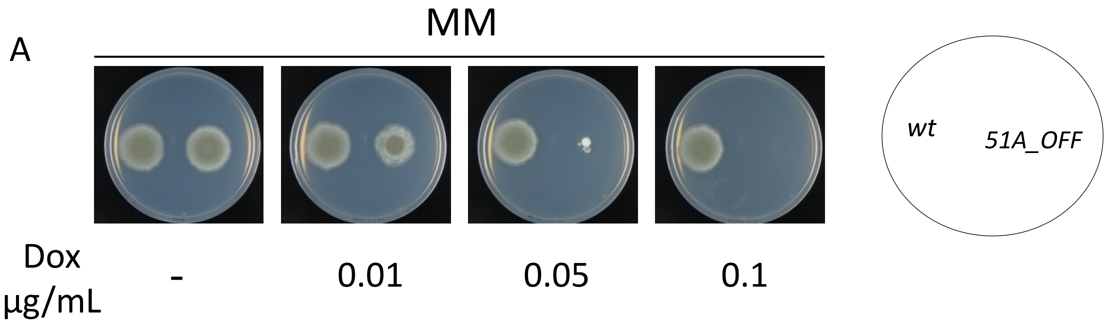

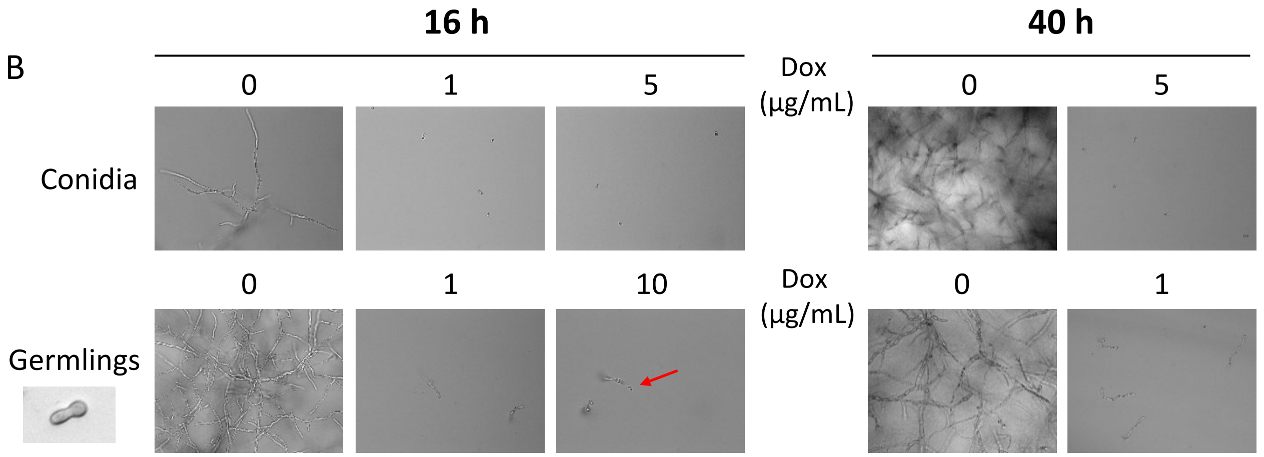

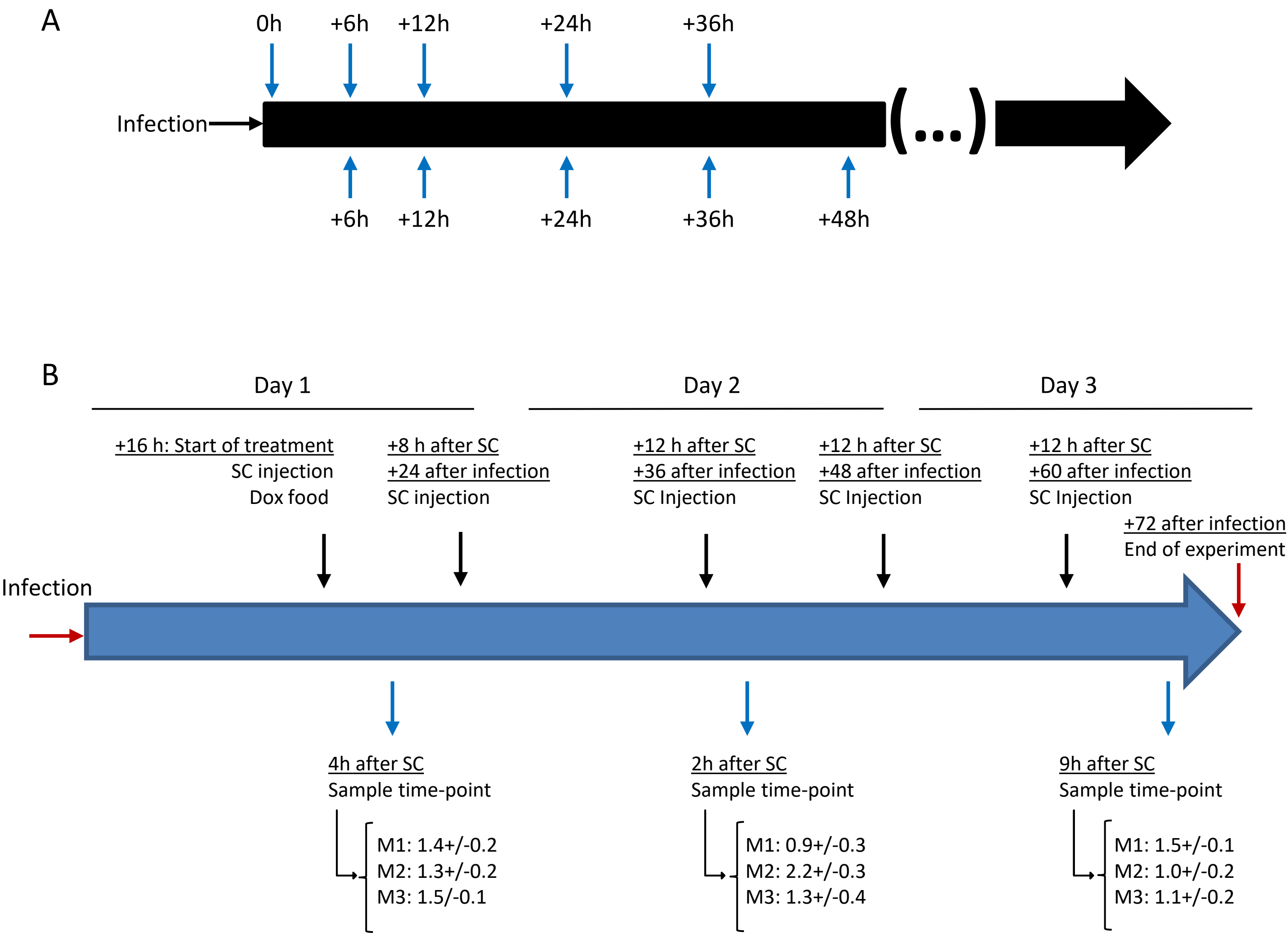
