## Supplementary material for "Targeting methionine synthase in a fungal pathogen causes a metabolic imbalance that impacts cell energetics, growth and virulence": Table S1

| Number metabolite | f. value | p. value | # | FDR | ID |
|---|---|---|---|---|---|
| 1 | 21,388 | 1,15E-08 | 7,9394 | 2,62E-08 | NA |
| 2 | 5,594 | 0,00157 | 2,8042 | 0,0021085 | Acetic acid, aminooxy- (3TMS) |
| 3 | 44,72 | 1,15E-12 | 11,939 | 4,23E-12 | NA |
| 4 | 54,449 | 7,87E-14 | 13,104 | 3,37E-13 | NA |
| 5 | 7,9211 | 0,000148 | 3,8298 | 0,0002186 | NA |
| 6 | 11,68 | 5,84E-06 | 5,234 | 1,01E-05 | NA |
| 7 | 26 | 1,17E-09 | 8,9325 | 2,95E-09 | Homoserine (2TMS) |
| 8 | 14,425 | 7,77E-07 | 6,1096 | 1,48E-06 | NA |
| 9 | 87,913 | 8,60E-17 | 16,066 | 5,77E-16 | Hydrogen sulfide (2TMS) |
| 10 | 14,816 | 5,95E-07 | 6,2258 | 1,14E-06 | Carbodiimide (2TMS) |
| 11 | 31,282 | 1,20E-10 | 9,9197 | 3,57E-10 | NA |
| 13 | 35,971 | 2,03E-11 | 10,692 | 6,29E-11 | NA |
| 15 | 23,185 | 4,54E-09 | 8,3432 | 1,08E-08 | Glyoxylic acid (1MEOX) (1TMS) |
| 18 | 5,1069 | 0,002684 | 2,5712 | 0,0035393 | Dodecane |
| 19 | 37,451 | 1,20E-11 | 10,92 | 3,90E-11 | Proline (1TMS) |
| 20 | 99,317 | 1,44E-17 | 16,842 | 1,09E-16 | Alanine, beta- (1TMS) |
| 21 | 14,834 | 5,87E-07 | 6,2311 | 1,14E-06 | Pyridine, 2-hydroxy- (1TMS) |
| 22 | 36,043 | 1,98E-11 | 10,704 | 6,18E-11 | Pyruvic acid (1MEOX) (1TMS) |
| 23 | 11,341 | 7,62E-06 | 5,118 | 1,29E-05 | Lactic acid (2TMS) |
| 24 | 4,237 | 0,007281 | 2,1378 | 0,0092577 | Dodecane |
| 25 | 7,9058 | 0,00015 | 3,8235 | 0,0002209 | Dodecane |
| 26 | 4,1472 | 0,008095 | 2,0918 | 0,010256 | Dodecane |
| 28 | 18,375 | 6,23E-08 | 7,2052 | 1,35E-07 | Pyruvic acid (2TMS) |
| 29 | 22,565 | 6,21E-09 | 8,2066 | 1,44E-08 | Valine (1TMS) |
| 30 | 26,041 | 1,15E-09 | 8,9406 | 2,92E-09 | Putrescine, N-methyl- (3TMS) |
| 31 | 55,472 | 6,08E-14 | 13,216 | 2,74E-13 | Sarcosine (2TMS) |
| 33 | 4,9922 | 0,003053 | 2,5153 | 0,0040102 | Dodecane |
| 34 | 6,0651 | 0,000948 | 3,0232 | 0,0012981 | Dodecane |
| 36 | 63,369 | 9,45E-15 | 14,025 | 4,80E-14 | Isovaleric acid, 2-oxo- (1MEOX) (1TMS |
| 37 | 23,316 | 4,25E-09 | 8,3719 | 1,02E-08 | Hydroxylamine (3TMS) |
| 38 | 22,735 | 5,70E-09 | 8,2442 | 1,33E-08 | NA |
| 42 | 3,3896 | 0,020222 | 1,6942 | 0,02457 | NA |
| 44 | 8,2996 | 0,000104 | 3,9841 | 0,0001565 | NA |
| 46 | 3,4306 | 0,019228 | 1,7161 | 0,023442 | NA |
| 47 | 17,918 | 8,19E-08 | 7,0868 | 1,76E-07 | NA |
| 48 | 8,7857 | 6,64E-05 | 4,1776 | 0,0001019 | Carbonic acid (1MEOX) (2TMS) |
| 49 | 7,9988 | 0,000137 | 3,8617 | 0,0002044 | Carbonic acid (1MEOX) (2TMS) |
| 50 | 542,88 | 6,56E-29 | 28,183 | 1,17E-26 | NA |
| 51 | 349,84 | 6,48E-26 | 25,188 | 3,84E-24 | NA |
| 52 | 67,53 | 3,84E-15 | 14,416 | 2,07E-14 | NA |
| 54 | 50,212 | 2,40E-13 | 12,621 | 9,69E-13 | Ethanolamine (3TMS) |
| 55 | 14,331 | 8,29E-07 | 6,0813 | 1,57E-06 | NA |
| 56 | 13,952 | 1,08E-06 | 5,9661 | 2,02E-06 | Unknown#bth-pae-013 |
| 57 | 3,1285 | 0,027967 | 1,5534 | 0,033636 | NA |
| 58 | 11,004 | 9,98E-06 | 5,0008 | 1,68E-05 | NA |
| 59 | 281,46 | 1,92E-24 | 23,717 | 8,54E-23 | NA |

| | | | | | |
|---|---|---|---|---|---|
| 61 | 9,6911 | 2,99E-05 | 4,525 | 4,70E-05 | NA |
| 62 | 131,95 | 2,09E-19 | 18,68 | 2,19E-18 | NA |
| 63 | 57,077 | 4,09E-14 | 13,388 | 1,92E-13 | NA |
| 64 | 22,736 | 5,69E-09 | 8,2445 | 1,33E-08 | Isoleucine (1TMS) |
| 65 | 5,4279 | 0,001881 | 2,7255 | 0,0025086 | NA |
| 68 | 8,7796 | 6,68E-05 | 4,1752 | 0,0001021 | NA |
| 70 | 41,622 | 3,00E-12 | 11,522 | 1,02E-11 | NA |
| 73 | 7,0974 | 0,000329 | 3,4825 | 0,000467 | NA |
| 76 | 4,982 | 0,003088 | 2,5103 | 0,0040417 | Valine (2TMS) |
| 77 | 25,986 | 1,18E-09 | 8,9296 | 2,95E-09 | NA |
| 78 | 69,704 | 2,45E-15 | 14,612 | 1,38E-14 | NA |
| 79 | 238,89 | 2,43E-23 | 22,614 | 6,67E-22 | NA |
| 80 | 167,48 | 5,67E-21 | 20,247 | 7,76E-20 | NA |
| 81 | 22,653 | 5,94E-09 | 8,2261 | 1,38E-08 | Butanoic acid, 4-hydroxy- (2TMS) |
| 82 | 69,401 | 2,60E-15 | 14,584 | 1,45E-14 | NA |
| 83 | 56,543 | 4,66E-14 | 13,332 | 2,15E-13 | Unknown#sst-cgl-008 |
| 84 | 40,069 | 4,97E-12 | 11,304 | 1,65E-11 | 1,3-Dihydroxyaceton (1MEOX) (2TMS |
| 86 | 46,562 | 6,68E-13 | 12,175 | 2,59E-12 | Serine (2TMS) |
| 87 | 46,303 | 7,21E-13 | 12,142 | 2,76E-12 | Serine (2TMS) |
| 88 | 44,556 | 1,21E-12 | 11,917 | 4,40E-12 | Alanine (2TMS) |
| 92 | 4,2739 | 0,006973 | 2,1566 | 0,0088969 | Alkane |
| 93 | 18,825 | 4,79E-08 | 7,3199 | 1,05E-07 | Phosphoric acid (3TMS) |
| 94 | 2,8483 | 0,039788 | 1,4002 | 0,047373 | Phosphoric acid (3TMS) |
| 95 | 20,155 | 2,25E-08 | 7,6481 | 5,07E-08 | Phosphoric acid (3TMS) |
| 96 | 4,5169 | 0,005252 | 2,2797 | 0,0067743 | NA |
| 101 | 9,4132 | 3,80E-05 | 4,4201 | 5,96E-05 | Threonine (2TMS) |
| 103 | 18,172 | 7,03E-08 | 7,1529 | 1,52E-07 | similar to Cyclohexasiloxane, dodecan |
| 105 | 8,2778 | 0,000106 | 3,9753 | 0,000159 | NA |
| 107 | 6,5015 | 0,000601 | 3,2208 | 0,0008332 | NA |
| 110 | 30,865 | 1,42E-10 | 9,8466 | 4,12E-10 | Succinic acid (2TMS) |
| 112 | 5,3933 | 0,001954 | 2,709 | 0,0025961 | NA |
| 113 | 258,45 | 7,20E-24 | 23,142 | 2,80E-22 | Glyceric acid (3TMS) |
| 114 | 220,55 | 8,35E-23 | 22,078 | 1,75E-21 | NA |
| 115 | 7,4496 | 0,000233 | 3,633 | 0,0003342 | Alkane |
| 119 | 58,534 | 2,88E-14 | 13,541 | 1,37E-13 | NA |
| 120 | 53,473 | 1,01E-13 | 12,996 | 4,23E-13 | NA |
| 121 | 37,837 | 1,05E-11 | 10,978 | 3,44E-11 | NA |
| 122 | 16,616 | 1,83E-07 | 6,7382 | 3,78E-07 | Alanine (3TMS) |
| 123 | 172,12 | 3,74E-21 | 20,428 | 5,54E-20 | NA |
| 124 | 229,05 | 4,66E-23 | 22,332 | 1,19E-21 | Mevalonic acid-1,5-lactone (1TMS) |
| 125 | 56,028 | 5,29E-14 | 13,276 | 2,42E-13 | NA |
| 126 | 74,328 | 9,76E-16 | 15,011 | 5,70E-15 | NA |
| 127 | 4,0579 | 0,009 | 2,0458 | 0,011321 | Tryptamine, 5-hydroxy- (4TMS) |
| 128 | 12,572 | 2,95E-06 | 5,5306 | 5,25E-06 | Alanine [+CO2] (2TMS) |
| 130 | 43,317 | 1,77E-12 | 11,753 | 6,16E-12 | NA |
| 131 | 14,086 | 9,84E-07 | 6,0071 | 1,84E-06 | Aspartic acid (2TMS) |
| 133 | 120,62 | 8,03E-19 | 18,095 | 7,33E-18 | Alanine, beta- (3TMS) |

| | | | | | |
|---|---|---|---|---|---|
| 134 | 96,471 | 2,21E-17 | 16,656 | 1,60E-16 | NA |
| 135 | 87,989 | 8,49E-17 | 16,071 | 5,77E-16 | NA |
| 136 | 158,26 | 1,34E-20 | 19,873 | 1,70E-19 | Tyrosine, 3-iodo- (3TMS) |
| 138 | 13,157 | 1,91E-06 | 5,7184 | 3,47E-06 | NA |
| 140 | 113,08 | 2,10E-18 | 17,677 | 1,83E-17 | NA |
| 141 | 30,784 | 1,47E-10 | 9,8323 | 4,22E-10 | NA |
| 142 | 54,683 | 7,41E-14 | 13,13 | 3,26E-13 | NA |
| 143 | 121,72 | 7,01E-19 | 18,154 | 6,57E-18 | NA |
| 144 | 43,996 | 1,43E-12 | 11,844 | 5,16E-12 | Malic acid (3TMS) |
| 147 | 180,1 | 1,87E-21 | 20,728 | 3,03E-20 | Threitol (4TMS) |
| 148 | 180,6 | 1,79E-21 | 20,746 | 3,03E-20 | NA |
| 149 | 16,456 | 2,02E-07 | 6,6941 | 4,14E-07 | Alkane |
| 150 | 6,8157 | 0,000437 | 3,3599 | 0,000612 | NA |
| 154 | 16,425 | 2,06E-07 | 6,6856 | 4,20E-07 | Pyroglutamic acid (1TMS) |
| 155 | 24,758 | 2,10E-09 | 8,6787 | 5,15E-09 | Pyroglutamic acid (1TMS) |
| 157 | 4,0902 | 0,008661 | 2,0624 | 0,010934 | Butanoic acid, 4-amino- (3TMS) |
| 158 | 12,822 | 2,45E-06 | 5,6115 | 4,42E-06 | NA |
| 159 | 3,6839 | 0,014107 | 1,8506 | 0,017498 | Glutamic acid (2TMS) |
| 160 | 3,0674 | 0,030191 | 1,5201 | 0,036068 | Glutamic acid (2TMS) |
| 161 | 6,0481 | 0,000965 | 3,0154 | 0,0013164 | NA |
| 163 | 9,7763 | 2,77E-05 | 4,5568 | 4,39E-05 | NA |
| 166 | 10,827 | 1,15E-05 | 4,9383 | 1,91E-05 | NA |
| 168 | 176,78 | 2,48E-21 | 20,605 | 3,84E-20 | NA |
| 169 | 22,362 | 6,90E-09 | 8,1612 | 1,58E-08 | NA |
| 172 | 32,575 | 7,22E-11 | 10,141 | 2,18E-10 | Alkane |
| 173 | 31,11 | 1,29E-10 | 9,8896 | 3,76E-10 | Phenylalanine (1TMS) |
| 174 | 49,943 | 2,58E-13 | 12,589 | 1,03E-12 | NA |
| 175 | 601,32 | 1,31E-29 | 28,883 | 4,66E-27 | NA |
| 176 | 43,833 | 1,51E-12 | 11,822 | 5,36E-12 | Glutaric acid, 2-hydroxy- (3TMS) |
| 177 | 221,9 | 7,60E-23 | 22,119 | 1,69E-21 | NA |
| 181 | 134,58 | 1,55E-19 | 18,808 | 1,74E-18 | NA |
| 182 | 219,22 | 9,16E-23 | 22,038 | 1,81E-21 | Glutaric acid, 2-oxo- (1MEOX) (2TMS) |
| 183 | 11,782 | 5,39E-06 | 5,2686 | 9,40E-06 | NA |
| 185 | 51,877 | 1,53E-13 | 12,815 | 6,27E-13 | NA |
| 187 | 65,114 | 6,44E-15 | 14,191 | 3,37E-14 | NA |
| 188 | 68,029 | 3,46E-15 | 14,461 | 1,89E-14 | NA |
| 189 | 12,591 | 2,91E-06 | 5,5368 | 5,20E-06 | NA |
| 191 | 113,6 | 1,97E-18 | 17,707 | 1,75E-17 | NA |
| 194 | 42,91 | 2,00E-12 | 11,698 | 6,92E-12 | Tartaric acid (4TMS) |
| 195 | 38,447 | 8,55E-12 | 11,068 | 2,82E-11 | NA |
| 196 | 7,8286 | 0,000162 | 3,7916 | 0,0002348 | NA |
| 197 | 7,6713 | 0,000188 | 3,7262 | 0,0002708 | NA |
| 199 | 7,7526 | 0,000174 | 3,76 | 0,0002515 | Benzoic acid, 4-hydroxy- (2TMS) |
| 200 | 6,9077 | 0,000398 | 3,4002 | 0,0005622 | Unknown#sst-cgl-044 |
| 201 | 7,8476 | 0,000159 | 3,7995 | 0,0002315 | Gluconic acid-1,4-lactone (4TMS) |
| 202 | 58,606 | 2,83E-14 | 13,548 | 1,36E-13 | NA |
| 204 | 309,16 | 4,46E-25 | 24,351 | 2,27E-23 | Xylose (1MEOX) (4TMS) BP |

| | | | | | |
|---|---|---|---|---|---|
| 205 | 27,268 | 6,57E-10 | 9,1825 | 1,72E-09 | Arabitol (5TMS) |
| 206 | 89,064 | 7,11E-17 | 16,148 | 4,96E-16 | NA |
| 208 | 109,38 | 3,45E-18 | 17,463 | 2,92E-17 | Arabitol (5TMS) |
| 209 | 17,288 | 1,20E-07 | 6,9203 | 2,53E-07 | Galactitol (6TMS) |
| 211 | 6,0649 | 0,000948 | 3,0232 | 0,0012981 | Alkane |
| 212 | 145,75 | 4,66E-20 | 19,331 | 5,54E-19 | NA |
| 214 | 19,653 | 2,98E-08 | 7,5261 | 6,58E-08 | NA |
| 215 | 74,576 | 9,30E-16 | 15,031 | 5,52E-15 | NA |
| 216 | 65,402 | 6,05E-15 | 14,218 | 3,21E-14 | NA |
| 217 | 9,0808 | 5,10E-05 | 4,2927 | 7,89E-05 | NA |
| 218 | 134,54 | 1,56E-19 | 18,807 | 1,74E-18 | Glycerol-3-phosphate (4TMS) |
| 219 | 28,599 | 3,67E-10 | 9,4357 | 9,81E-10 | Alkane |
| 220 | 132,54 | 1,95E-19 | 18,709 | 2,11E-18 | NA |
| 221 | 28,314 | 4,15E-10 | 9,3823 | 1,09E-09 | Ornithine (3TMS) |
| 222 | 5,5705 | 0,00161 | 2,7931 | 0,0021549 | NA |
| 224 | 25,819 | 1,27E-09 | 8,8961 | 3,16E-09 | NA |
| 225 | 5,8918 | 0,001139 | 2,9434 | 0,001542 | NA |
| 226 | 60,143 | 1,97E-14 | 13,706 | 9,74E-14 | similar to Fructose Derivate |
| 227 | 16,553 | 1,90E-07 | 6,7208 | 3,91E-07 | NA |
| 228 | 7,3738 | 0,000251 | 3,6008 | 0,0003585 | similar to Fructose Derivate |
| 229 | 6,0219 | 0,000992 | 3,0034 | 0,0013482 | NA (Sugar) |
| 234 | 10,727 | 1,25E-05 | 4,903 | 2,05E-05 | NA |
| 235 | 125,47 | 4,45E-19 | 18,351 | 4,31E-18 | NA |
| 237 | 36,593 | 1,63E-11 | 10,789 | 5,17E-11 | NA (Sugar) |
| 238 | 64,47 | 7,41E-15 | 14,13 | 3,82E-14 | Glyceric acid-3-phosphate (4TMS) |
| 239 | 108,58 | 3,85E-18 | 17,415 | 3,11E-17 | Citric acid (4TMS) |
| 240 | 384,3 | 1,49E-26 | 25,826 | 1,33E-24 | NA |
| 242 | 17,256 | 1,23E-07 | 6,9116 | 2,57E-07 | NA |
| 243 | 10,807 | 1,17E-05 | 4,9312 | 1,93E-05 | NA |
| 244 | 33,076 | 5,95E-11 | 10,225 | 1,81E-10 | Galactose_1_5TMS |
| 245 | 30,392 | 1,73E-10 | 9,7627 | 4,84E-10 | NA |
| 246 | 28,516 | 3,80E-10 | 9,4201 | 1,01E-09 | Galactose_1_5TMS |
| 247 | 4,4744 | 0,005517 | 2,2583 | 0,0070907 | Altrose (1MEOX) (5TMS) BP |
| 248 | 3,8933 | 0,010956 | 1,9604 | 0,013733 | Psicose (1MEOX) (5TMS) BP |
| 249 | 3,1627 | 0,026798 | 1,5719 | 0,03245 | Allose (1MEOX) (5TMS) MP |
| 251 | 17,317 | 1,18E-07 | 6,9279 | 2,50E-07 | Sorbose (1MEOX) (5TMS) MP |
| 252 | 14,916 | 5,56E-07 | 6,2553 | 1,08E-06 | Fructose (1MEOX) (5TMS) MP |
| 253 | 8,8132 | 6,48E-05 | 4,1885 | 9,99E-05 | Ribose (1MEOX) (4TMS) BP |
| 254 | 10,273 | 1,82E-05 | 4,7396 | 2,91E-05 | Ribose (1MEOX) (4TMS) BP |
| 255 | 10,883 | 1,10E-05 | 4,9583 | 1,84E-05 | NA |
| 257 | 11,534 | 6,54E-06 | 5,1842 | 1,13E-05 | NA |
| 258 | 24,888 | 1,97E-09 | 8,7056 | 4,87E-09 | NA |
| 261 | 6,763 | 0,000461 | 3,3367 | 0,0006429 | NA (Sugar) |
| 266 | 17,809 | 8,75E-08 | 7,0582 | 1,86E-07 | NA (Sugar) |
| 268 | 26,262 | 1,04E-09 | 8,9849 | 2,67E-09 | NA (Sugar) |
| 271 | 5,1242 | 0,002633 | 2,5796 | 0,0034843 | Tyrosine (2TMS) |
| 274 | 10,604 | 1,38E-05 | 4,859 | 2,25E-05 | Galactose (1MEOX) (5TMS) MP |

| | | | | | |
|---|---|---|---|---|---|
| 275 | 15,537 | 3,67E-07 | 6,4355 | 7,30E-07 | Mannitol (6TMS) |
| 281 | 3,745 | 0,013099 | 1,8828 | 0,016304 | Alkane |
| 282 | 20,574 | 1,78E-08 | 7,7484 | 4,05E-08 | NA |
| 284 | 3,4897 | 0,01788 | 1,7476 | 0,021949 | NA |
| 288 | 8,4924 | 8,68E-05 | 4,0615 | 0,0001315 | Pentadecanoic acid, 14-methyl-, meth |
| 291 | 201,53 | 3,34E-22 | 21,476 | 5,95E-21 | NA |
| 294 | 4,8448 | 0,003606 | 2,4429 | 0,0046882 | NA426001 |
| 295 | 15,036 | 5,12E-07 | 6,2906 | 1,01E-06 | NA426001 |
| 296 | 10,87 | 1,11E-05 | 4,9535 | 1,85E-05 | Glucopyranose [-H20] (4TMS) |
| 298 | 78,78 | 4,22E-16 | 15,374 | 2,64E-15 | NA |
| 299 | 11,466 | 6,90E-06 | 5,1609 | 1,19E-05 | Glucopyranose, D- (5TMS) |
| 300 | 13,167 | 1,90E-06 | 5,7214 | 3,47E-06 | NA |
| 302 | 4,8443 | 0,003608 | 2,4427 | 0,0046882 | Galactose_1_5TMS |
| 303 | 3,07 | 0,030091 | 1,5216 | 0,036068 | Galactose_1_5TMS |
| 305 | 6,1906 | 0,000831 | 3,0806 | 0,0011462 | Gluconic acid-1,4-lactone (4TMS) |
| 306 | 3,7592 | 0,012876 | 1,8902 | 0,016083 | NA (Sugar) |
| 311 | 12,777 | 2,53E-06 | 5,5969 | 4,55E-06 | NA |
| 312 | 6,8522 | 0,000421 | 3,3759 | 0,0005922 | similar to NA (Inositol like) |
| 314 | 355,21 | 5,11E-26 | 25,292 | 3,64E-24 | NA |
| 316 | 109,09 | 3,59E-18 | 17,445 | 2,97E-17 | Hexadecenoic-acid_2_1TMS |
| 319 | 45,613 | 8,83E-13 | 12,054 | 3,31E-12 | NA |
| 320 | 34,249 | 3,82E-11 | 10,418 | 1,17E-10 | Hexadecanoic acid (1TMS) |
| 321 | 13,211 | 1,84E-06 | 5,7356 | 3,37E-06 | NA |
| 322 | 125,42 | 4,48E-19 | 18,349 | 4,31E-18 | Inositol, myo- (6TMS) |
| 323 | 166,34 | 6,29E-21 | 20,202 | 8,29E-20 | Xylulose-5-phosphate (1MEOX) (5TMS |
| 324 | 256,99 | 7,87E-24 | 23,104 | 2,80E-22 | NA (Sugar) |
| 325 | 8,2345 | 0,00011 | 3,9578 | 0,0001648 | NA |
| 326 | 10,599 | 1,39E-05 | 4,8573 | 2,25E-05 | NA (Sugar) |
| 327 | 7,8905 | 0,000152 | 3,8172 | 0,0002232 | NA (Sugar) |
| 328 | 4,3681 | 0,006244 | 2,2045 | 0,0079959 | NA |
| 329 | 81,033 | 2,81E-16 | 15,551 | 1,79E-15 | NA |
| 330 | 52,65 | 1,25E-13 | 12,903 | 5,18E-13 | Octadecadienoic acid methyl ester, 9, |
| 331 | 203,08 | 2,97E-22 | 21,527 | 5,57E-21 | NA |
| 332 | 29,913 | 2,10E-10 | 9,6769 | 5,76E-10 | NA |
| 333 | 15,898 | 2,90E-07 | 6,5382 | 5,86E-07 | NA |
| 335 | 48,882 | 3,46E-13 | 12,461 | 1,37E-12 | NA |
| 336 | 28,995 | 3,10E-10 | 9,5093 | 8,41E-10 | Octadecatrienoic acid methylester, 9, |
| 337 | 241,14 | 2,11E-23 | 22,677 | 6,67E-22 | NA |
| 338 | 55,261 | 6,41E-14 | 13,193 | 2,85E-13 | Octadecanoic acid methyl ester (FAM |
| 340 | 10,656 | 1,33E-05 | 4,8776 | 2,16E-05 | Heptadecanoic acid (1TMS) |
| 341 | 8,7032 | 7,16E-05 | 4,1452 | 0,0001089 | NA |
| 343 | 45,784 | 8,39E-13 | 12,076 | 3,18E-12 | NA |
| 344 | 9,8076 | 2,70E-05 | 4,5685 | 4,29E-05 | NA |
| 345 | 43,793 | 1,53E-12 | 11,817 | 5,38E-12 | Glycerophosphoglycerol (5TMS) |
| 346 | 105,55 | 5,85E-18 | 17,233 | 4,63E-17 | NA |
| 347 | 85,883 | 1,21E-16 | 15,918 | 7,82E-16 | NA |
| 348 | 19,373 | 3,49E-08 | 7,457 | 7,67E-08 | NA |

| | | | | | |
|---|---|---|---|---|---|
| 349 | 30,228 | 1,85E-10 | 9,7335 | 5,14E-10 | NA |
| 350 | 58,853 | 2,67E-14 | 13,574 | 1,30E-13 | Gluconic acid (6TMS) |
| 351 | 170,47 | 4,33E-21 | 20,364 | 6,16E-20 | Mannitol (6TMS) |
| 352 | 101,91 | 9,83E-18 | 17,007 | 7,61E-17 | NA |
| 353 | 15,538 | 3,67E-07 | 6,4359 | 7,30E-07 | NA |
| 355 | 3,6706 | 0,014336 | 1,8436 | 0,017721 | Ornithine, N2-acetyl- (4TMS) |
| 358 | 3,4563 | 0,018628 | 1,7298 | 0,022789 | Glycerol-3-phosphate (4TMS) |
| 359 | 48,388 | 3,97E-13 | 12,401 | 1,55E-12 | Octadecadienoic acid, n- (1TMS) |
| 360 | 26,157 | 1,09E-09 | 8,9639 | 2,78E-09 | Octadecenoic acid, 9-(E)- (1TMS) |
| 361 | 98,053 | 1,74E-17 | 16,76 | 1,29E-16 | NA |
| 364 | 44,994 | 1,06E-12 | 11,974 | 3,93E-12 | NA |
| 365 | 54,138 | 8,51E-14 | 13,07 | 3,61E-13 | NA |
| 366 | 2,8067 | 0,041938 | 1,3774 | 0,049766 | NA |
| 368 | 14,125 | 9,57E-07 | 6,0189 | 1,80E-06 | NA |
| 369 | 70,835 | 1,94E-15 | 14,711 | 1,12E-14 | Octadecanoic acid (1TMS) |
| 371 | 42,257 | 2,46E-12 | 11,61 | 8,41E-12 | NA |
| 372 | 153,96 | 2,03E-20 | 19,691 | 2,50E-19 | NA |
| 373 | 28,735 | 3,46E-10 | 9,4611 | 9,33E-10 | NA |
| 374 | 7,9962 | 0,000138 | 3,8607 | 0,0002044 | Mannose-6-phosphate (1MEOX) (6TM |
| 375 | 7,1224 | 0,000321 | 3,4933 | 0,0004573 | Xylulose-5-phosphate (1MEOX) (5TMS |
| 377 | 5,8743 | 0,001161 | 2,9353 | 0,0015652 | NA (Sugar-phosphate) |
| 379 | 24,11 | 2,87E-09 | 8,5424 | 6,95E-09 | NA |
| 380 | 24,208 | 2,73E-09 | 8,5632 | 6,67E-09 | NA (Sugar-phosphate) |
| 381 | 6,6492 | 0,000517 | 3,2865 | 0,000719 | NA (Sugar-phosphate) |
| 382 | 54,48 | 7,80E-14 | 13,108 | 3,37E-13 | NA |
| 383 | 76,532 | 6,41E-16 | 15,193 | 3,87E-15 | NA |
| 384 | 15,633 | 3,44E-07 | 6,4631 | 6,92E-07 | NA |
| 385 | 76,835 | 6,06E-16 | 15,218 | 3,72E-15 | NA |
| 386 | 240,02 | 2,26E-23 | 22,645 | 6,67E-22 | NA |
| 387 | 86,769 | 1,04E-16 | 15,983 | 6,86E-16 | NA |
| 389 | 445,56 | 1,47E-27 | 26,834 | 1,74E-25 | NA |
| 390 | 223,46 | 6,82E-23 | 22,166 | 1,62E-21 | NA |
| 391 | 14,683 | 6,51E-07 | 6,1865 | 1,25E-06 | Inositol-2-phosphate, myo- (7TMS) |
| 392 | 11,264 | 8,11E-06 | 5,0912 | 1,37E-05 | NA |
| 393 | 61,42 | 1,47E-14 | 13,834 | 7,35E-14 | NA |
| 396 | 10,588 | 1,40E-05 | 4,8534 | 2,26E-05 | NA |
| 397 | 4,5804 | 0,004881 | 2,3115 | 0,0063181 | NA |
| 398 | 14,943 | 5,45E-07 | 6,2633 | 1,07E-06 | NA |
| 400 | 30,645 | 1,56E-10 | 9,8077 | 4,40E-10 | NA |
| 402 | 12,166 | 4,01E-06 | 5,3969 | 7,03E-06 | NA |
| 404 | 9,4016 | 3,84E-05 | 4,4157 | 5,98E-05 | NA |
| 406 | 92,474 | 4,10E-17 | 16,387 | 2,92E-16 | NA |
| 408 | 36,495 | 1,68E-11 | 10,774 | 5,31E-11 | NA |
| 414 | 12,396 | 3,37E-06 | 5,4728 | 5,96E-06 | NA |
| 415 | 16,852 | 1,58E-07 | 6,8026 | 3,28E-07 | NA |
| 416 | 126,11 | 4,13E-19 | 18,384 | 4,20E-18 | NA (Sugar) |
| 417 | 13,359 | 1,65E-06 | 5,7821 | 3,06E-06 | NA |

| | | | | | |
|---|---|---|---|---|---|
| 418 | 9,3991 | 3,85E-05 | 4,4147 | 5,98E-05 | NA |
| 419 | 13,248 | 1,79E-06 | 5,7472 | 3,30E-06 | NA |
| 420 | 19,703 | 2,90E-08 | 7,5382 | 6,44E-08 | NA |
| 422 | 11,395 | 7,30E-06 | 5,1366 | 1,25E-05 | Kestose, 6- (11TMS) |
| 424 | 26,813 | 8,06E-10 | 9,0939 | 2,09E-09 | NA |
| 425 | 11,378 | 7,40E-06 | 5,1307 | 1,26E-05 | NA |
| 426 | 37,027 | 1,40E-11 | 10,855 | 4,48E-11 | NA |
| 427 | 30,717 | 1,51E-10 | 9,8204 | 4,31E-10 | Adenosine (3TMS) (Derivate not found |
| 433 | 29,947 | 2,07E-10 | 9,683 | 5,73E-10 | NA |
| 434 | 23,193 | 4,52E-09 | 8,345 | 1,08E-08 | NA |
| 438 | 31,971 | 9,15E-11 | 10,039 | 2,74E-10 | NA |
| 439 | 31,109 | 1,29E-10 | 9,8894 | 3,76E-10 | NA |
| 441 | 15,527 | 3,69E-07 | 6,4327 | 7,30E-07 | NA |
| 442 | 10,569 | 1,42E-05 | 4,8467 | 2,28E-05 | NA |
| 447 | 20,071 | 2,36E-08 | 7,6278 | 5,27E-08 | NA |
| 451 | 40,692 | 4,05E-12 | 11,392 | 1,36E-11 | NA |
| 453 | 12,3 | 3,62E-06 | 5,4414 | 6,38E-06 | NA |
| 454 | 3,1541 | 0,027086 | 1,5673 | 0,032687 | NA |
| 455 | 3,5824 | 0,015962 | 1,7969 | 0,019663 | NA |

12-(Z,Z)-, n-

12,15-(Z,Z,Z)-, n-
